# Growth in confinement promotes *Pseudomonas aeruginosa* tolerance to antibiotics

**DOI:** 10.1101/2025.10.15.682596

**Authors:** Sourabh Monnappa, Zainebe Al-Mayyah, Mahmut Selman Sakar, Alexandre Persat

## Abstract

Bacteria often proliferate within confined spaces imposed by host tissues, extracellular matrices, or their own biofilms. In such environments, cells press against surrounding materials and experience elevated mechanical stress, but whether these forces influence pathogen physiology and fitness remains unclear. Here, we show that *Pseudomonas aeruginosa* adapts to mechanical confinement by increasing resilience to antibiotics. Using synthetic hydrogels of tunable stiffness that restrict physical expansion without limiting nutrient access, we demonstrate that growth in elastic materials reduces *P. aeruginosa* sensitivity to multiple clinically relevant antibiotics in a stiffness-dependent manner. Although slower growth contributes to this decreased susceptibility, Tn-seq under antibiotic treatment identified key regulators of mechanically induced tolerance. We find that active efflux mediated by sodium– proton *Sha* antiporters, together with protective remodeling of the bacterial membrane, enhances the resilience of confined populations without impacting growth. These findings reveal that *P. aeruginosa* adapts to mechanical stress in ways that may promote treatment failure even in the absence of intrinsic antibiotic resistance.

## Introduction

During the course of host colonization or in the wild, bacteria experience mechanical forces as they interact with abiotic and biological materials^1–4^. These forces are now being recognized as important regulators of bacterial physiology and pathogenicity, particularly in the context of surface adaptation^1,5,6^. As they transition between planktonic and sessile lifestyles, bacteria sense surface contact to deploy their virulence arsenal, reinforce adhesion, or initiate biofilm formation. *Pseudomonas aeruginosa*, for example, uses a mechanosensory systems to detect surface association^7^, leading to the upregulation of its type III secretion system and timely deployment of its twitching motility machinery^8,9^.

Bacteria also colonize the bulk of biological solid materials when they grow within abscesses, form lesions between cells, infect intracellularly, or elongate within their own biofilm matrix^5,10–15^. As they expand, single bacteria deform their elastic surrounding and in turn experience a restoring mechanical stress referred to as growth-induced pressure^1,16,17^. These forces can passively impact the organization of single cells in the confined community. For example *Vibrio cholerae* embedded within agarose hydrogel grows into oblate ellipsoidal clusters whose internal ordering arises from confinement-driven alignment^18^. In addition, *V. cholerae* and *P. aeruginosa* growth generate internal stress within their own biofilms, which can lead to buckling instabilities that deform underlying materials, including epithelial tissues^19^.

While mechanics explain the cellular arrangement of collectives in confinement, how growth-induced pressure impacts bacterial physiology remains poorly understood. Investigating mechanobiology of microbes in the bulk of elastic materials requires simultaneous physiological monitoring and mechanical stimulation with careful control of chemical perturbation. In *Escherichia coli*, growth to saturation in microfluidic chambers upregulates extracellular matrix components, suggesting that growth-induced stress can trigger biofilm formation^20^. Transcriptomic profiling of *E. coli* encapsulated in methacrylated alginate further revealed that stiff hydrogels limit growth by impairing nutrient uptake and metabolic activity^21,22^. While these results suggest that bacteria respond to mechanical confinement, whether growth induced pressure or starvation cause these phenotypic changes remains unclear. A recent investigation leveraging a microfluidic system that decouples mechanics and nutrient depletion showed that *E. coli* responds to confinement by activating the *Rcs* pathways to adjust growth and cell division^23^, indicating that bacteria can actively adapt to pressure. Still, despite its ubiquity during infection, the functional impact and adaptation mechanisms to confinement remain largely underexplored.

*P. aeruginosa*, a prominent opportunistic pathogen that causes acute and chronic infections^24^, represents an ideal model to investigate how confined growth shapes bacterial physiology. *P. aeruginosa* can replicate within airway and corneal epithelial cells^25,26^, lesions, abscesses^27,28^ and extracellular matrix of infected tissues^29,30^, thereby exposing populations to mechanical confinement. In the lungs of cystic fibrosis patients, *P. aeruginosa* grows within thick, viscoelastic mucus, which restricts movement and spreading^24,31^. Finally, it very commonly forms antibiotic tolerant biofilms where single cells surround themselves with a self-secreted elastic matrix^32–35^. Antibiotic treatment failure against *P. aeruginosa* is often attributed to genetic resistance, but tolerance also plays a substantial role, as many chronic isolates lack resistance determinants^35,36^. With its multitude of efflux pumps, *P. aeruginosa* displays intrinsic tolerance to diverse antimicrobials^37–39^, which is further amplified within biofilms, where effective minimum inhibitory concentrations (MIC) can increase by orders of magnitude^38,40–42^. However, our current understanding of regulation of tolerance has largely focused on chemical regulation of efflux systems, while the role of mechanics has been overlooked. We here hypothesize that *P. aeruginosa* enhances antibiotic tolerance in response to mechanical confinement. Using synthetic hydrogels with tunable mechanical properties, we specifically investigate how physical constraints influence *P. aeruginosa* survival under antibiotic exposure.

## Results

### Confinement enhances *P. aeruginosa* tolerance to colistin

To confine *P. aeruginosa* without affecting nutrient access, we cross-linked synthetic, mechanically-tunable polyethylene glycol (PEG) hydrogels around single cells^43–46^. After mixing prepolymer with a bacterial inoculum and initiating crosslinking, PEG forms a structurally homogenous elastic network with high water content (> 99% by weight) that uniformly immobilizes single bacteria (**Figure 1A and Figure S1A**). The crosslinking process did not affect *P. aeruginosa* viability (**Figure S1D**). Upon medium supplementation, single cells proliferated within the hydrogel to form three-dimensional multicellular bacterial clusters organized in tightly-packed communities of contiguous cells **(Figure 1B)**. These clusters thus experience growth-induced pressure generated by elastic deformations of the hydrogel.

**Figure 1.**
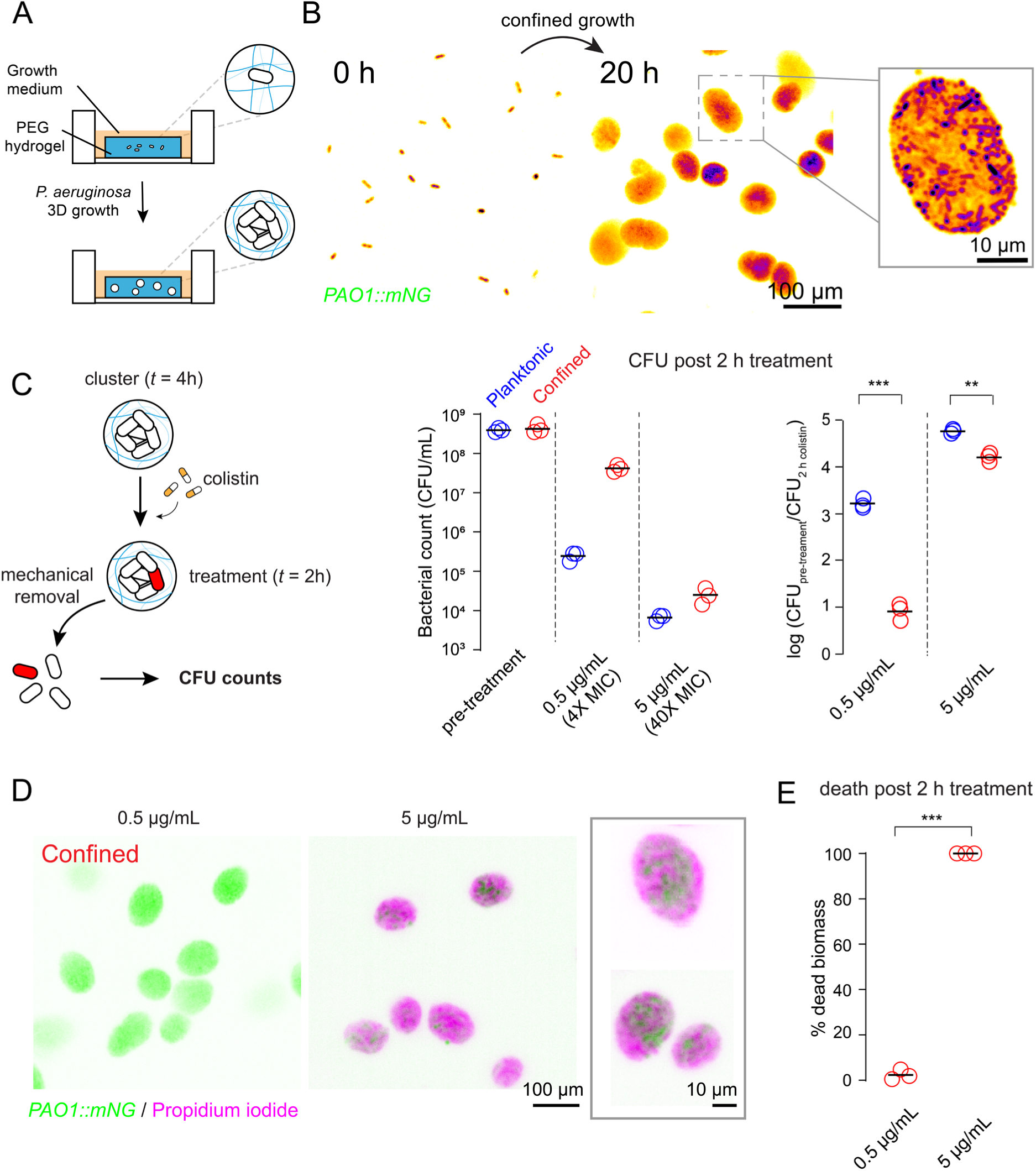
**Confinement increases *P. aeruginosa* tolerance to colistin**. **(A)** Schematic illustration of the experimental setup used to confine *P. aeruginosa* within PEG hydrogels. **(B)** Confocal fluorescence microscopy images showing isolated *P. aeruginosa* constitutively expressing *mNeonGreen* at 0 h and clusters at 20 h. The boxed region shows a magnified view captured (Inset). **(C**, left) Schematic of colistin treatment of confined *P. aeruginosa*. Cell viability assessment after 2-h colistin treatment by CFU counts for both planktonic (middle, blue circles) and confined cells (red circles) with computed log fold reduction (right). Data point: biological replicate. Black line: mean biological triplicates. Two-tailed Welch’s *t*-test (\*\*\**p* = 0.00023 at 4× MIC and \*\*\**p* = 0.0034 at 40× MIC). **(D)** Confocal images of confined *PAO1*::*mNeonGree*n in 1 kPa hydrogel treated with colistin and stained with propidium iodide. Live cells are shown in green, while dead cells are in magenta. Inset shows a zoomed-in view highlighting spatial patterns of cell death. **(E)** Dead biomass quantification of bacterial clusters at two different colistin concentrations (for > 100 clusters in each biological triplicates).

We next tested how confinement influenced *P. aeruginosa* antibiotic sensitivity by challenging clusters to colistin, a clinically-relevant last resort antibiotic^47^. After 4 h of growth, we treated small clusters with colistin at 4 and 40 times its MIC. We first assessed sensitivity relative to planktonic bacteria by liberating cells from confinement after 2 h of treatment and plating them for colony-forming unit (CFU) (**Figure 1C**). While colistin was highly effective against planktonic populations in liquid culture, treatment in confined conditions showed higher survival. At 4x MIC, the number of confined cells decreased only 10-fold compared to a 1000- fold killing for planktonic populations (**Figure 1C**). At 40x MIC (5 μg/mL), CFU measurements showed a stronger reduction in overall bacterial population in both populations, but confined cells still survived significantly better than planktonic cells (**Figure 1C**).

A physical mechanism wherein antibiotics diffuse relatively poorly through the hydrogel could cause the observed loss of antibiotic sensitivity^20^. To test this possibility, we quantified how PEG hydrogel influenced the diffusion of small molecules modeled by FITC-dextran (MW 10,000) using fluorescence recovery after photobleaching (FRAP). We measured no significant differences in the diffusion coefficient of FITC-dextran between hydrogel and free solution, indicating that PEG does not impact molecular diffusion of small molecules (**Figure S2A-C**). We also wondered whether transport within the clusters could affect drug penetration as in biofilms. Reduced penetration tends to initially cause cell death at the outer edge of the population while sparing the core. To test this, we imaged spatial patterns of cell death in the clusters upon antibiotic exposure by staining with propidium iodide (PI) after 2 h of exposure to colistin. At 0.5 μg/mL, most planktonic cells stained positive for PI, indicating those had compromised membranes^48^, with roughly 60% of the population showing cell death (**Figure S3**). By contrast, only a few bacteria in clusters were PI positive, consistent with CFU quantification (**Figure 1D**). To further exhibit patterns of cell death, we increased colistin concentration. We could not detect live cells in planktonic populations, while a small fraction of cells in confined clusters survived (**Figure 1E and Figure S3**). We could not distinguish any specific pattern of cells deaths as PI-negative cells were visible both at the edge and in the core of clusters (**Figure 1D**), indicating uniform penetration of colistin. Altogether, our results indicate that *P. aeruginosa* growing in physically constrained environments tolerates colistin via a mechanism independent of molecular transport.

### Tolerance of confined clusters increases with hydrogel stiffness

To explore a mechanism wherein confined growth mechanically stimulates *P. aeruginosa* tolerance, we sought to modulate the magnitude of the growth-induced pressure. Clusters experience an elastic force that scales with the Young’s modulus of the hydrogel and their size^49–54^. To modulate pressure independently of cluster size, we produced soft (1 kPa, as above) and stiff (23 kPa) PEG hydrogels for cluster formation (**Figure 2A and Figure S1B**). These two hydrogel compositions have marginal differences in pore size (**Figure S1C**), and FRAP measurements confirmed that they did not cause detectable changes in diffusion of FITC-dextran compared to liquid conditions (**Figure S2C**). We verified antibiotic penetration in clusters grown in stiff hydrogels using rhodamine B–labelled polymyxin B, a fluorescent analog of colistin (also known as polymyxin E). Imaging showed rapid antibiotic penetration within 5 min in both soft and stiff environments (**Figure S5**). Based on a simple physical model accounting for the size of the cavities and the stiffness of the hydrogel^55^, we calculated that *P. aeruginosa* clusters experience about 0.6 kPa growth induced pressure in soft hydrogels and 14.2 kPa in stiff ones after 4 h of growth, the time of colistin treatment (**Figure S4A,C**).

**Figure 2.**
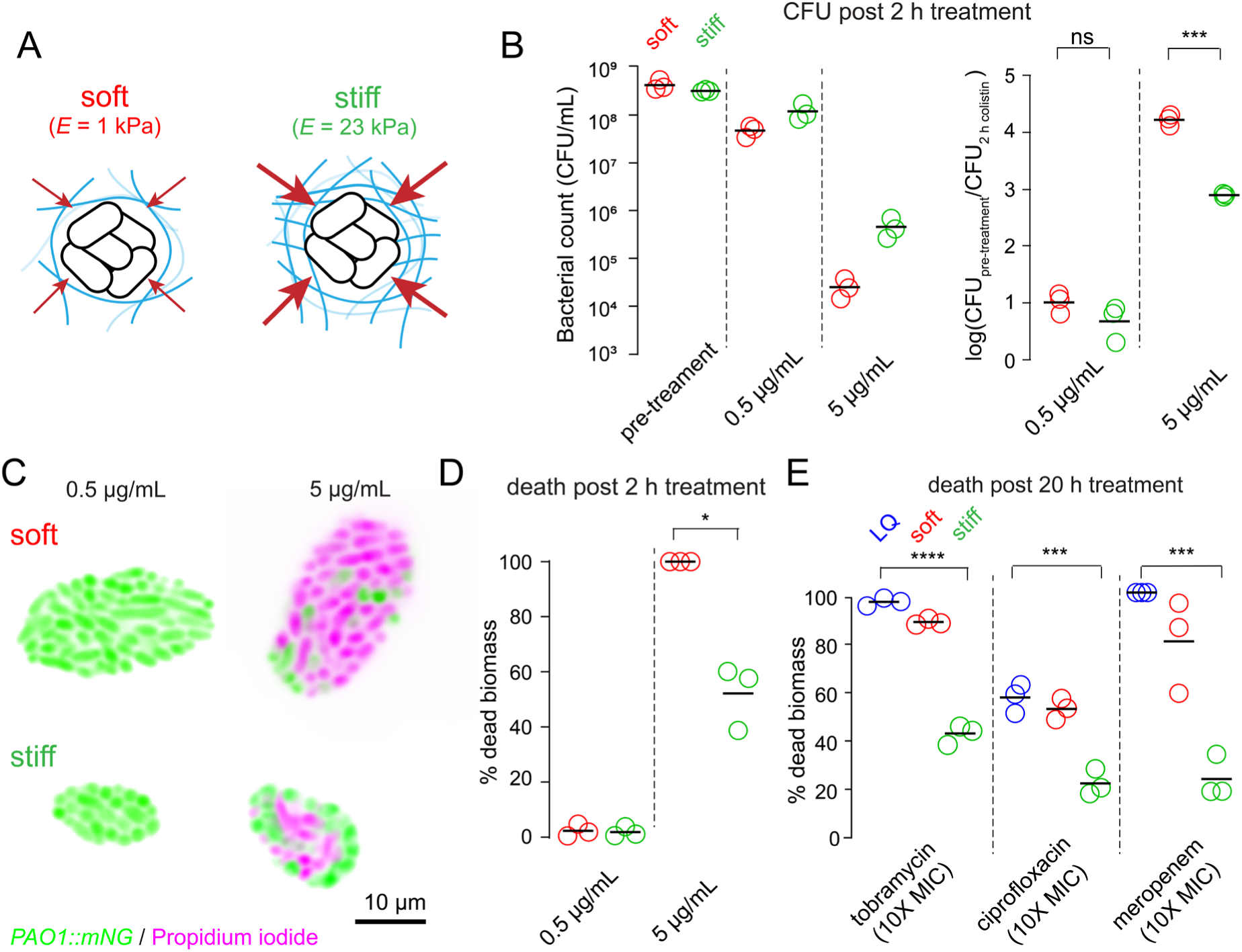
**Hydrogel mechanical stiffness enhances tolerance to multiple antibiotic classes**. **(A)** Illustration of compressive forces acting on clusters within PEG hydrogels with two different stiffness and mesh density. **(B)** Survival measurements after colistin exposure using CFU counts (left). Log-reduction plots (right) indicate fold-change in survival to colistin treatment (Welch’s unpaired two-sided *t*-test, \*\*\**p* = 0.00080). **(C)** High-resolution 100x confocal images (cross-sectional view) of colistin- treated *P*. *aeruginosa* clusters clusters in 1 kPa and 23 kPa conditions at two colistin concentrations. Live cells are in green, while dead cells are in magenta. **(D)** Quantification of percentage dead biomass in *P. aeruginosa* clusters. For 5 µg/ml colistin, soft vs stiff populations differ significantly (Welch’s unpaired two-sided *t*-test, \**p* = 0.0193). Scale bar is 10 μm. **(E)** Quantification of dead biomass of planktonic (LQ), soft and stiff hydrogel clusters, after 20 h treatment with tobramycin (10 μg/mL), ciprofloxacin (1.25 μg/mL), and meropenem (60 μg/mL). Tukey’s MSD test showed significant differences between planktonic and stiff hydrogel for tobramycin (\*\*\*\**p* < 0.0001), ciprofloxacin (\*\*\**p* = 0.0004), and meropenem (\*\*\**p* = 0.0006).

We next measured their survival after colistin exposure by CFU counting. At 0.5 μg/mL, clusters in both conditions remained largely viable after 2 h of exposure. At 5 μg/mL, cells in the stiff hydrogel survived 10-fold better than the ones in the soft hydrogels, suggesting that increased pressure significantly improves survival (**Figure 2B**). At 0.5 μg/mL, there was no detectable PI staining in clusters from either stiffness, confirming minimal cell death (**Figure 2C**). However, at a higher dose (5 μg/mL), dead biomass increased significantly in soft hydrogel whereas clusters within 23 kPa retained a substantial fraction of live cells (**Figure 2C**). Quantitative analysis showed that approximately 50% of the bacterial population survived in the stiff matrix compared to near-complete loss of viability in the soft matrix (**Figure 2D**), confirming CFU measurements.

Given the apparent dose-dependent response, we measured colistin minimum inhibitory concentration (MIC) in confined conditions by imaging cluster growth when exposed to antibiotics right after encapsulation (**Figure S6A**). Confinement strikingly increased colistin tolerance, with cells growing at 2x MIC in soft and 4x MIC in stiff hydrogels (**Figure S6B,C**), reinforcing that the survival advantage arises from substrate mechanics rather than impaired drug accessibility. In conclusion, mechanically confined growth of *P. aeruginosa* promote antibiotic tolerance, highlighting how physical constraints can influence bacterial survival under antibiotic stress.

Colistin disrupts the outer membrane integrity by binding to the lipid A component of lipopolysaccharides^56,57^, suggesting mechanical confinement promotes tolerance by altering membrane properties. Before investigating specific mechanisms, we tested whether confinement influences sensitivity to antibiotics with alternative mechanisms of action^56^, the aminoglycoside tobramycin, the fluoroquinolone ciprofloxacin, and the β-lactam meropenem. We again combined CFU and imaging to evaluate *P. aeruginosa* viability under each treatment. Across all tested antibiotics, survival was consistently higher in confined conditions relative to planktonic populations after 20 h of treatment, with enhanced tolerance in stiff hydrogels than in soft ones (**Figure 2E, Figure S8**). Even after shorter treatments, cells growing in stiff matrix survived 100- to 1000-times better than planktonic cells (**Figure S7A-C, i**). We further examined single-cell antibiotic responses under confinement using high-resolution imaging 5 h after treatment, followed by PI staining. Ciprofloxacin-treated cells exhibited diverse morphologies, including filamentation, and intracellular puncta suggestive of stalled replication complexes^58,59^ (**Figure S7B, ii**). Meropenem induced shape changes, but these appeared less severe under confinement, even in cells that lost viability (**Figure S7C, ii**). These results show that confinement and hydrogel stiffness enhance antibiotic tolerance and support *P. aeruginosa* survival through a pleiotropic mechanisms.

### Mechanical confinement slows bacterial growth

Given the broad tolerance spectrum, we first envisioned a simple mechanism wherein mechanics reduces drug sensitivity by impacting growth rate^57,59,60^. We investigated *P. aeruginosa* growth behavior under confinement by imaging cluster proliferation at the single- cell level (**Figure 3A, ii**). Volume quantification over time shows that clusters reach a maximum size at about 20 h of growth (**Figure 3A, i and Figure 3B**). We compared growth across hydrogel stiffness from 1 kPa to 63 kPa to cover a wide range of mechanical properties representative of soft biological materials^45,46^. Imaging those clusters after 6 h of growth showed pronounced differences in cluster size with increasing stiffness (**Figure 3C**). Time course experiments demonstrated that the cluster growth rate was decreasing with increasing stiffness even at early times (**Figure 3D, i**). After 20 h, cluster volume in stiffer gels was about 5-fold lower than in the softest gel (**Figure 3D, ii**). The differences in cluster size and volumetric expansion between hydrogel stiffnesses suggest a change in growth state which could shape *P. aeruginosa* antibiotic sensitivity.

**Figure 3.**
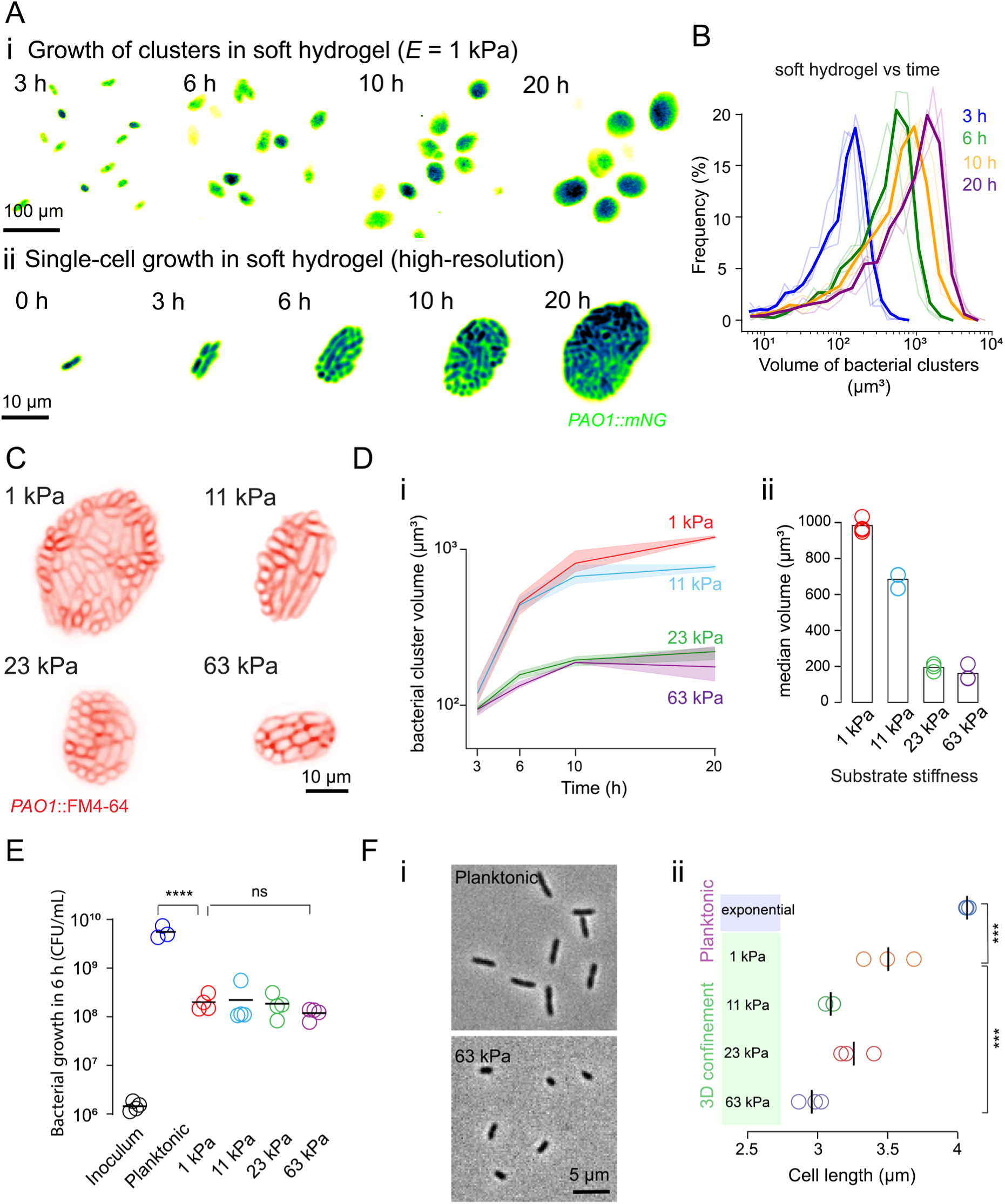
Confinement mechanically constrains *P. aeruginosa* growth. (**A, i**) Time-lapse images of *PAO1-mNeonGreen* growth within 1 kPa PEG hydrogels. (**A, ii**) High-resolution scanning confocal images of single-cell growing into a cluster over time. (**B**) Cluster volume distributions in 1 kPa PEG hydrogel over time. (**C**) Confocal fluorescence images of PAO1 clusters in hydrogels of different stiffness labeled with FM4-64 after 6 h of growth. (**D, i**) Volumetric growth of *PAO1-mNG* clusters under varying PEG hydrogel stiffness. The solid thick line shows the mean of the biological triplicates, while the shaded region represents the standard deviation across replicates. (**D,ii**) Bar graph comparing the median cluster volume at 20 h of growth in hydrogels (bar: mean of biological triplicates). (**E**) Number of cells after 6 h of growth in hydrogels. Solid circle: single biological replicate, black line: mean of biological replicates (*N* = 4). Statistics: one-way ANOVA between planktonic vs 1 kPa (\*\*\*\**p* = 0.0001), while 1 kPa vs 63 kPa (*p* > 0.99). (**F, i**) Phase contrast images of single-cells from planktonic and 63 kPa confined conditions grown for 6 h. (**F, ii**) Cell length measurements after 6 h in hydrogels across stiffness. Circles: the mean cell length for each biological replicate. One-way ANOVA (*p* < 0.0001), Tukey’s HSD test between log-phase and 1 kPa (\*\*\**p* = 0.0005), and 1 kPa and 63 kPa (\*\*\**p* = 0.0006).

Hydrogel stiffness impacts growth induced pressure buildup (**Figure S4C**), which could slow down bacterial division and/or elongation rate. To further explore these mechanisms, we measured the number of cell divisions per unit time with increasing stiffness. Confined cells showed approximately a 1.5-log fold reduction in CFUs compared to planktonic cultures, demonstrating that confinement decreases division rate (**Figure 3E**). We then further asked whether division decreased with increasing growth-induced pressure. CFU counts remained similar across material properties, showing that increased substrate stiffness does not affect division rate. A live/dead assay to determine whether cell death contributed to the observed cluster size differences showed little difference across stiffnesses (**Figure S9**).

Next, we stained bacterial membranes with FM4-64 and imaged individual cells within clusters to directly measure their size. In situ imaging showed that cells were larger in the softest hydrogel (**Figure 3C and Figure S10A, i**). Since cell orientation within clusters posed a challenge to accurate cell size quantification, we extracted cells from the hydrogel for imaging **(Figure 3F, i and Figure S10B)**. Confined *P. aeruginosa* populations were significantly shorter

compared to planktonic cells, with cells in 63 kPa hydrogels being the shortest (**Figure 3F, ii and Figure S10A, ii**). Together, these results demonstrate that mechanical confinement decreases bacterial elongation rate in a stiffness-dependent manner, and negatively impacts replication rate independently of stiffness. Such mechanism would naturally impact tolerance to growth-dependent antibiotics such as ciprofloxacin, but in less obvious ways for others. We therefore speculate that growth-induced pressure drives cells into a physiological state with broader effects on physiology that could be independent of growth. We therefore further explored mechanisms of tolerance by investigating the physiology of cells in confinement.

### Cells released from confinement resume growth but maintain tolerance

To test whether mechanical adaptation had lasting effects, we investigated the tolerance of freshly-released bacteria to measure their growth rate and drug sensitivity. We first confined *P. aeruginosa* during their active growth phase (4 h), released them from hydrogels as planktonic cells which were immediately transferred to fresh medium for live microscopic imaging (**Figure 4A)**. After cell segmentation, we quantified the rate of change of the normalized cell surface area for the first 30 min to measure growth (**Figure 4B**). We could not distinguish differences in growth rate between planktonic and previously-confined cells, regardless of hydrogel stiffness (**Figure 4C**). This shows that mechanical confinement does not impair the ability of cells to resume normal growth once released. Therefore, if tolerance of confined cells was only due to growth defects, cells released from the hydrogels should become as sensitive as planktonic cells.

**Figure 4.**
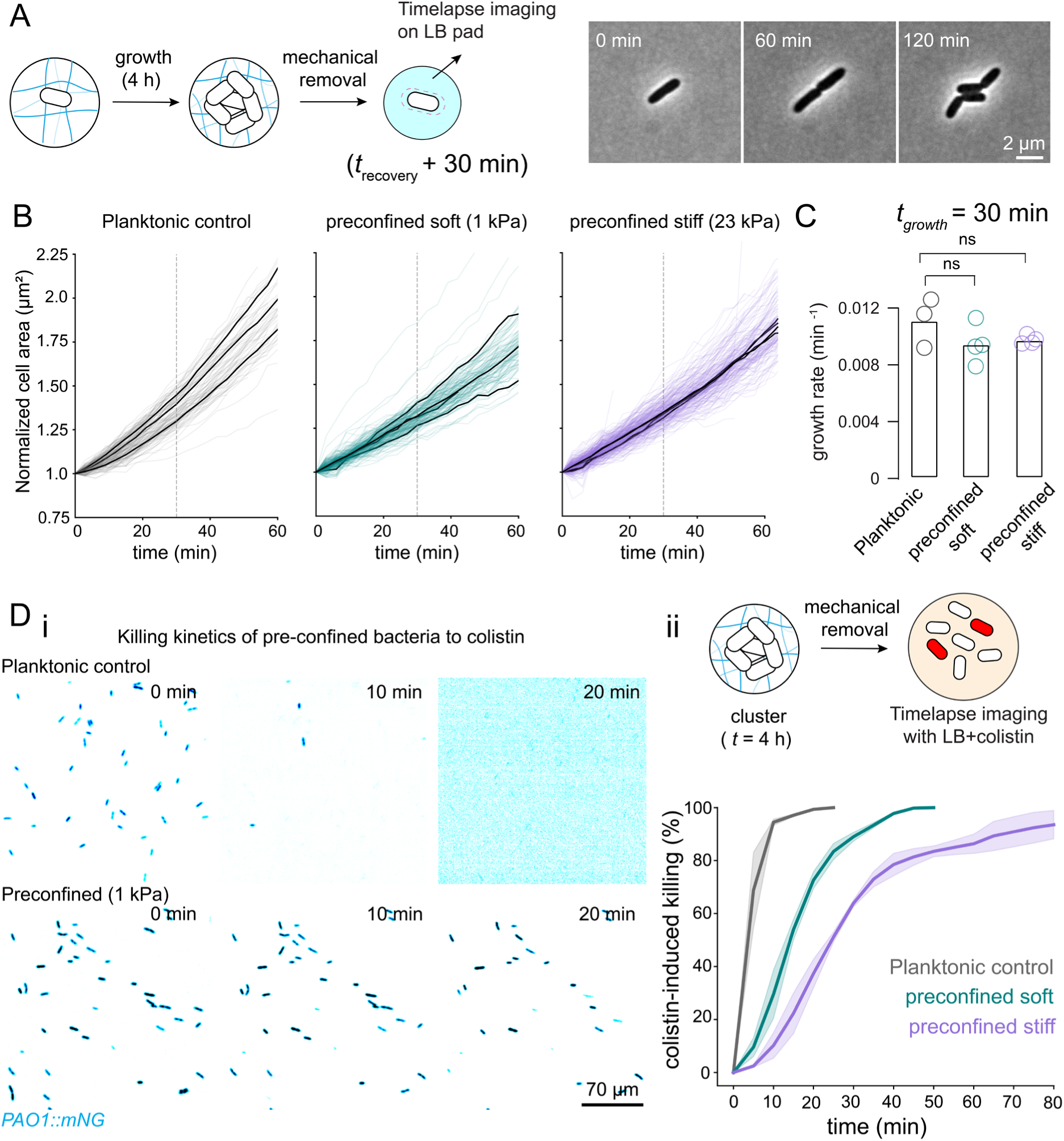
Released bacteria recover growth but maintain colistin tolerance. **(A)** Schematic summary of regrowth experiments. Representative phase-contrast images showing released PAO1 growth monitored by live microscopy. **(B)** Normalized cell area over time in the regrowth assay for planktonic cells as control, cells released from 1 kPa gels and 23 kPa gels. Each fine colored line represents the growth of a single cell out of the biological triplicate (solid black lines). **(C)** Growth rates of planktonic and released *P. aeruginosa* cells computed from the first 30 min. Bars represent the mean growth rate for each condition; individual biological replicates are shown as overlaid colored circles. Statistics: unpaired two-tailed *t*-test, Mann–Whitney, *p* = 0.4 for soft; *p* = 0.6268 for stiff. **(D, i)** Fluorescence images capturing the impact of colistin on *PAO1:mNeonGreen* after release. **(D, ii)** Graphical outline of the killing kinetics assay. Killing curves of planktonic and confined PAO1 after exposure to colistin. The solid thick line indicates the mean of the biological replicates, while the shaded region depicts the standard deviation across replicates. Statistical analysis compared the time to 50% cell death across all conditions. Pairwise Welch’s *t*-tests: liquid vs soft: *p* = 0.003; liquid vs stiff: *p* = 0.0004; soft vs stiff: *p* < ***).

We thus tested whether effects of confinement still had an impact on antibiotic sensitivity after release by immediately exposing them to colistin during the microscopy growth assay. We dynamically monitored antibiotic efficacy by quantifying the loss of fluorescence signal from *mNeonGreen* expressing PAO1 cells. In the planktonic cell control, we observed lysis of 80% of the population within 9 min of colistin exposure (**Figure 4D, i-top and 4D, ii**). By contrast, cells previously confined in 1 kPa hydrogel survived colistin treatment for a longer time, as only 80% of the population had lysed after 24 min of exposure (**Figure 4D, i-below and 4D, ii**). Killing was further delayed in cells previously-confined within stiffer hydrogels, as 80% of the population was lysed only after 43 min of exposure **(Figure 4D, ii)**. Released cells also showed enhanced tolerance to ciprofloxacin despite their similar growth rates (**Figure S11**). This suggests that mechanical stress induces an adaptive response to confinement whose effects extend beyond growth rate changes.

### A functional-genomic screen links membrane remodeling to tolerance

To decode the molecular mechanisms of confinement-induced tolerance, we implemented a functional genomics screen of populations under antibiotic selection in hydrogels. Using transposon sequencing (Tn-seq), we searched mutations that impact antibiotic sensitivity in confined conditions relative to planktonic growth^61^. We first grew a PAO1 Tn5-based transposon library^62^ in stiff hydrogels for 4 h and sequenced the resulting population to identify mutations altering growth in confinement relative to liquid cultures^63^ (See Material and methods). Tn-seq identified 74 genes (**Supplementary Table 5, i**) whose mutations impacted *P. aeruginosa* fitness under confinement compared to planktonic reference (45 with decreased fitness and 29 with increased fitness). Depleted mutants included genes involved in cell division and lipopolysaccharide biosynthesis, while enriched mutants included several two-component systems (**Supplementary Table 5, iv**). These first results show that cells in confinement have distinct fitness requirements compared to free-living cells. (**Figure S12A)**.

We next performed Tn-seq under colistin and tobramycin exposure for both planktonic and confined cultures (**Figure 5A)**. As before, we allowed cells to grow for 4 h in stiff hydrogels before introducing antibiotics, after which confined populations were sequenced to compare fitness to treatment in liquid culture (**Supplementary Table 5, ii and iii**). We found 112 insertions that increased and 52 that decreased fitness under colistin treatment, while under tobramycin exposure, 731 insertions decreased and 324 increased fitness. These results reveal broad shifts in genetic requirements when confined cells are exposed to antibiotics. We could not observe a strong correlation between fitness during growth in confinement and fitness under antibiotic exposure, consistent with our hypothesis that a growth alone does not explain mechanically-induced tolerance (**Figure S12B,C**).

**Figure 5.**
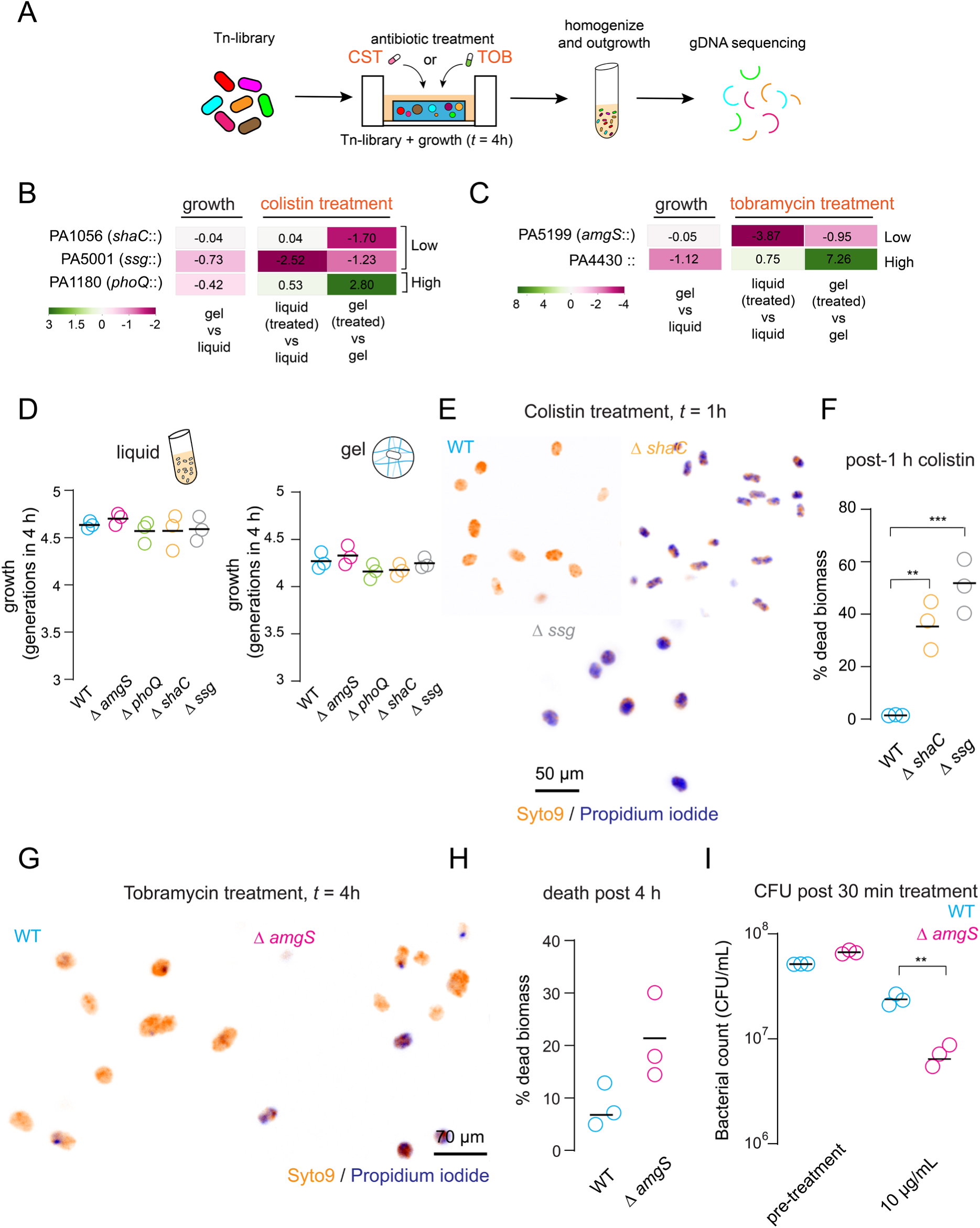
**A functional-genomic screen shows membrane remodeling stimulates antibiotic tolerance under confinement**. **(A)** Schematic representation Tn-seq in the presence of antibiotics. CST, colistin; TOB, tobramycin. See materials and methods for more details. Fitness effects of transposon insertions for selected genes under colistin (**B**) and tobramycin (**C**) treatment in gel. Genes passing the significance cut-off (adjusted *p* < 0.05, TRANSIT resampling) in antibiotic-treated gel were compared to growth in untreated gel and also to liquid culture treated with corresponding antibiotic. (See materials and methods and Supplementary table 5, iii). **(D)** CFU-based validation of deletion mutant growth and fitness in liquid culture (left) and stiff gel (right). Each data point represents an independent biological replicate (N = 3); horizontal black lines mark their means. Statistics: one-way ANOVA (*p* = 0.6112). **(E)** Live/dead staining of WT and deletion mutants grown as clusters in gel and treated with colistin for 1 h. Orange marks live cells, and blue highlights the dead population. **(F)** Quantification of percentage dead biomass in WT and mutant strains. Statistic: ANOVA for *shaC*: \*\**p* = 0.0035, *ssg:* \*\*\**p* = 0.0004. **(G)** Fluorescence images of tobramycin-treated clusters of WT and Δ*amgS* mutant. **(H)** Cell death quantification after 4 h of tobramycin treatment. **(I)** CFU count after 30 min tobramycin exposure for WT (blue circles) and Δ*amgS* (pink circles). Circle: biological replicate; black line means of biological triplicates. Statistics: unpaired *t*-test with Welch’s correction, \*\**p* = 0.0078).

Several mutations resensitized *P. aeruginosa* to colistin in confinement included lipopolysaccharide (LPS) related genes (*galU*, *ssg* (PA5001), and *wapH* (PA5004)). In addition, mutations in the two-component system genes *phoQ* increased fitness under colistin exposure, pointing to stress-sensing pathways that regulate membrane remodeling permitting antibiotic resilience^64^ (**Figure S12B and Figure S13B**; **See also Supplementary Table 5, iii**). Under tobramycin exposure, *mexZ* mutation increased the fitness of confined populations but to a lower extent than planktonic cells validating the role of efflux regulation in driving tolerance but also showing a milder effect in confined conditions. Disruptions in several electron transport chain genes also improved fitness, supporting the idea that tobramycin impacts bacterial metabolism under confinement (**Figure S12C and Figure S13D**; **See also Supplementary Table 5, ii**).

Next, we selected candidate genes that show significant fitness changes under antibiotic treatment. From these, we filtered genes whose mutations induced fitness changes in confined conditions only when exposed to antibiotics. Additionally, we included two genes that showed fitness defects both during confined growth and under drug exposure. The selected mutants were summarized as heatmaps with their fitness changes across tested experimental conditions. For colistin, we focused on genes involved in membrane homeostasis: *shaC*, *ssg* and *phoQ* (**Figure 5B**). For tobramycin, we included PA4430 and *amgS* as representative mutants (**Figure 5C**).

We validated the tolerance phenotypes for those genes with in-frame clean deletion mutants. In confinement, all mutants replicated to the same extent as WT as measured by CFU, confirming those genes are not essential to grow under pressure (**Figure 5D**). Although CFUs under confinement were unaffected, volume analysis showed that Δ*shaC* and Δ*ssg* formed slightly larger clusters than WT (1.5-fold and 1.2-fold larger, respectively; **Figure S14**). We next verified whether the Δ*ssg* and Δ*shaC* mutants reduced tolerance to colistin in confined conditions (5 μg/ml). Consistent with Tn-seq data, fluorescence imaging revealed a substantial increase in cell death in both mutants relative to WT (**Figure 5E**). Quantification of PI fluorescence showed colistin killed nearly 50% of the embedded population compared to only 2% of the WT population in clusters (**Figure 5F**). By contrast with planktonic behavior where it showed an expected increased sensitivity^64,65^, Δ*phoQ* showed a strong increase in tolerance under confinement. After 2 h of colistin exposure in stiff hydrogels, 92% of the phoQ population survived, compared to 51% of WT (**Figure S15AB**). Thus, both Δ*ssg* and Δ*shaC* mutants recover near planktonic level sensitivity to colistin when growing in confinement, Δ*phoQ* instead shows enhanced tolerance, without significant alteration in growth rate compared to WT.

We also ran a similar analysis for *amgS* deletion mutant under tobramycin treatment (10 μg/ml). *amgS* encodes a membrane-bound sensor kinase in the AmgRS two-component system, which regulates envelope stress and aminoglycoside tolerance^66^. PI staining showed increased cell death in the mutant (**Figure 5G**), and fluorescence quantification confirmed the higher proportion of non-viable cells (**Figure 5H**). A CFU-based survival assay after 30 min of drug exposure further showed an approximate 10-fold drop in viability for Δ*amgS* compared to WT (**Figure 5I**). Altogether, these results show that cells use cell envelope and membrane adaptation to better tolerate antibiotic exposure during confined growth.

## Discussion

Chemical signals such as starvation are often thought to increase tolerance, especially in the context of biofilms^32,33,36,67^. By decoupling molecular transport from mechanical stimulation, we showed that adaptation to growth induced pressure contributes to survival under antibiotic stress. Mechanically-induced tolerance may play a critical role in highly confined infection sites, such as abscesses^68^ or deep-seated infections, where treatment is problematic even in the absence of resistance and many times require surgical intervention^32,69^.

Confinement-induced stress influences growth in diverse organisms^51,70,71^. In our system, growth defects manifest themselves by a reduction in cell size. This is reminiscent of observations in *E. coli*, where growth-induced pressure slows growth and drives progressive size reduction through reductive division^23^, and in yeast^70^ where cell length decreases as stiffness increases. Upon release of pressure, cells recovered and rapidly resumed normal division^72^. By contrast, the effects of confinement on antibiotic tolerance perpetuated after release. This indicates an adaptation to growth in confinement that goes beyond forcing instantaneous effects on growth.

Confinement-driven remodeling of the lipopolysaccharide barrier forms the first barrier against colistin. Core lipopolysaccharide biosynthesis genes such as PA5001 (*ssg*), which encodes a cell-surface glycan protein^64,73,74^ may modulate membrane properties under compression to limit colistin binding. Consistent with this idea, *Δssg* mutants formed larger clusters and were more susceptible to colistin than WT, indicating that surface remodeling helps stabilize membrane in confined environments. Cells also rely on ion and membrane potential homeostasis to survive. Here, the sodium-proton (*Sha*) antiporter complex emerges as a critical player. This conserved transport system is known to balance sodium and proton flux to maintain pH and proton motive force^75,76^, both essential for overall electrochemical gradients. Mutations in *shaACD* were depleted under colistin treatment when confined. *ΔshaC* clusters were larger than WT, indicating a potential imbalance in osmotic pressure. We speculate that the *Sha* system could counter ion leakage during colistin induced perturbations, providing mechanochemical stabilization of the inner membrane. In doing so, these transport complex link ion-efflux to colistin tolerance. This previously unrecognized role highlights *Sha* antiporters as key determinants of antibiotic survival under confinement. Together with surface remodeling, these shift tolerance mechanisms away from lipid A modifications via PhoQ-dependent regulation^65,77,78^ which has a negative impact in confinement.

Antimicrobial treatment failure remains one of the most urgent challenges in global health, often driven by non-inherited tolerance rather than genetic resistance. Our findings reveal that bacteria within deep-seated infections, lesions, and abscesses may withstand antibiotic treatment through the mechanical effects of confinement. Understanding these biophysical mechanisms could inspire the development of new therapeutic strategies that target the regulators of mechanically induced tolerance. Together, our results establish mechanical confinement as a previously unrecognized determinant of antibiotic tolerance, highlighting the role of physical forces in shaping infection outcomes.

## Materials and Methods

### PEG hydrogel fabrication

To synthesize hydrogels, we prepared solutions of 4-arm poly(ethylene glycol) norbornene (PEG-NB) and 4-arm poly (ethylene glycol) thiol as precursor monomers, along with lithium phenyl-2,4,6-trimethylbenzoylphosphinate (LAP) as the photoinitiator in M9 minimal medium (without calcium and magnesium). Molecular weight (10,000 and 20,000 Da) and concentration (2.5% wt/vol, 5% wt/vol, and 15% wt/vol) of monomers to obtain hydrogels with different stiffnesses, while keeping the concentration of LAP constant at 2 mM. A mixture of monomers, photoinitiator and bacteria was done in the following ratios as described in supplementary Table 1, to control the stiffness of the hydrogel.

### Mechanical characterization of PEG hydrogels

#### **(i)** Bulk modulus

The bulk modulus of the hydrogel was determined by testing samples polymerized in cylindrical PDMS molds. After immersing these cylinders in M9 overnight, we analyzed them using a rheometer (TA Instruments) in compression mode at a deformation rate of 20 μm/s. Before testing, we measured the cylinder diameters with a digital caliper and defined their heights on the gap distance at which force began to increase. We obtained the elastic modulus from the slope of the linear fit of the stress-strain curves within the 15% strain range. The final Young modulus values calculated were averaged across the replicates. The fabricated hydrogels exhibited moduli ranging from 1 to 63 kPa, aligning with the bulk modulus of bacterial biofilm matrix, abscesses and lung tissues^46^.

#### **(ii)** Mesh size

The mesh size of the hydrogels was estimated using equilibrium swelling theory, following previously established protocols^79,80^. For each hydrogel cylinder, the volume and mass were measured in three different states: relaxed (r), swollen (s), and dry (d). The initial volume immediately after polymerization (Vr) and the volume after 24 hours of immersion in M9 (Vs) were determined using a digital caliper. To remove residual salts, the swollen hydrogels were washed in deionized water and subsequently dried overnight in an oven at 80°C. After drying, the mass of the dry network (Md) was recorded, and the dry volume (Vd) was calculated using the equation Vd = Md / ρPEG, where ρPEG was taken as 1.18 g/mL. The average molecular weight between cross-links (Mc) was determined using the below equation.

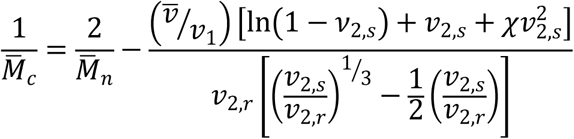

Here, v represents the specific volume of the polymer, which is 0.93 mL/g for PEG. V1 denotes the molar volume of water, taken as 18 mL/mol. The parameter corresponds to the polymer-solvent interaction parameter, with a value of 0.426 for PEG in water. The term M̅c refers to the average molecular weight of the polymer before cross-linking. Finally, 𝑣𝑣_2,𝑟𝑟_ and 𝑣𝑣_2,𝑠𝑠_ represent the polymer volume fractions in the relaxed and swollen states, respectively.

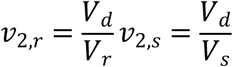

Finally, the mesh size (ξ) was calculated using,

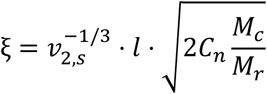

where l is the bond length along the polymer backbone (0.15 nm), Cn represents the Flory characteristic ratio (PEG = 4), and Mr corresponds to the molecular weight of the repeat unit (44 g/mol).

#### **(iii)** Diffusion coefficient using FRAP

To quantify the diffusion of molecules within PEG hydrogels, we used fluorescein isothiocyanate (FITC)-labelled dextran (10,000 MW) at a concentration of 0.5 mg mL⁻¹. PEG hydrogels with stiffnesses, 1 kPa and 23 kPa were polymerized and incubated overnight in the FITC-dextran solution at room temperature with gentle shaking (100 rpm). After incubation, hydrogels were washed twice with deionized water to remove unbound dextran. To visualize hydrogel positioning to FRAP, 2 µm red fluorescent beads were added to the prepolymer mix before polymerization. FRAP experiments were conducted on a confocal laser scanning microscope using a 20× air objective. A region of interest (ROI) of 16.6 × 16.6 pixels was selected within a 128 × 128-pixel field of view, with a pixel size of 1.84 µm. Photobleaching was performed using a 488 nm laser at 30% power for 5 sec, followed by acquisition at 250 msec intervals for a total of 30 sec. We measured the mean fluorescence intensity within the bleached ROI, and values were normalized to the pre-bleach intensity. A non-bleached internal control was also used Recovery curves were fitted using the non-linear exponential function using a custom Python script using scipy. optimize.curve_fit function:

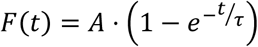

Where A is the maximum change in fluorescence intensity after bleaching, and τ is the time constant of fluorescence recovery. The half-time recovery (t1/2) was calculated as:

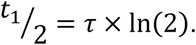

We then estimated the diffusion coefficient D using the standard two-dimensional FRAP diffusion equation:

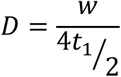

Where *w* is the radius of the bleached area in micrometres, and *t*1/2 is the time to half-maximal recovery. This equation assumes isotropic diffusion in 2D and is commonly used in FRAP analysis of hydrogels^81^.

### Bacterial growth in confinement

To prepare hydrogels, we flushed a prepolymer solution containing *P. aeruginosa* bacteria into a microfluidic channel made of PDMS (100 μm in height and 1 mm in length) on a Matek glass-bottom dish. We cleaned the glass dish with isopropanol and Milli-Q water, then functionalized with 3-(trimethoxysilyl)propyl methacrylate to covalently adhere the hydrogel to the dish. To achieve this, we washed for 10 minutes in a solution composed of 1 mL of 3-(trimethoxysilyl) propyl methacrylate, 6 mL of diluted acetic acid (1:10 acetic acid: water), and 200 mL of 70% ethanol. After washing, we rinsed with ethanol and dried in an oven at 60°C for 10 minutes.

Next, we added 8 μL of the prepolymer solution through the inlet of the channel and placed it under UV light (365 nm) for 1 minute to allow cross-linking. Immediately after polymerization, the PDMS channel was removed using tweezers and washed the hydrogel 2–3 times with M9 salt solution to eliminate unpolymerized monomers. Finally, addition of LB growth medium to the Matek dish and incubated at 37°C to allow the bacteria to grow and form 3D clusters.

### Bacterial strains and culture conditions

All the bacterial strains were grown in LB medium at 37°C, 225 rpm shaking (revolutions per minute). Overnight bacterial cultures were diluted 1:1000 in fresh LB and grown until the exponential phase (optical density at 600 nm: 0.5 - 0.6). Fluorescent PAO1-*mNG* was grown overnight in LB supplemented with gentamycin (60 µg/ml) and then diluted 1:100 in antibiotic-free LB and grown to exponential phase (OD600: 0.5-0.6). A full list of strains is provided in supplementary table 2.

### Bacterial strain construction

We used *Pseudomonas aeruginosa* strain PAO1^82^ as the genetic background for all experiments. Supplementary Table 2 lists all bacterial strains, plasmids, and primers used in this study. To construct unmarked, in-frame deletions, we first amplified ∼750 to 1,000 bp regions upstream and downstream of each target gene from PAO1 genomic DNA. After purifying the PCR products with the Monarch PCR & DNA Cleanup Kit (New England Biolabs), we assembled them into the EcoRI-HF and XbaI-digested pEX18Gm suicide vector ^83,84^ using Gibson assembly. We then transformed the resulting plasmids into *Escherichia coli* XL10-Gold (Agilent) and selected gentamicin-resistant colonies on LB agar (10 µg/ml gentamicin). We screened transformants by colony PCR and verified the assembled constructs by Sanger sequencing (Microsynth). Next, we extracted the verified plasmids using the GeneJET Plasmid Miniprep Kit (Thermo Scientific) and introduced them into *E. coli* S17^83^ via electroporation. We used this strain as a donor for biparental conjugation with PAO1, following previously described protocols^85^. After conjugation, we selected merodiploid colonies on Vogel–Bonner Minimal Medium containing 60 µg/ml gentamicin^85^. To isolate recombinants with successful deletions, we plated the merodiploids on LB without NaCl and supplemented with 10% sucrose. We then screened sucrose-resistant colonies by PCR and confirmed clean deletions by Sanger sequencing.

To generate strains for microscopy, we used published constructs encoding *mNeonGreen* under control of the constitutive *tet* promoter. We inserted these constructs at the *att*Tn7 site in the PAO1 genome by electroporation and selected gentamicin-resistant colonies on LB (60 µg/ml) ^31,85^.

### Antibiotic tolerance assay

#### **(i)** Colony forming units (CFU) method

Overnight PAO1 WT cultures were diluted 1:1000 in fresh LB and grown at 37°C with shaking at 225 rpm until reaching the exponential phase (optical density at 600 nm: 0.5–0.6). The culture was then centrifuged at 8500 rpm for 3 minutes, resuspended in 500 µL of M9 solution, and filtered using a 5.0 µm pore-size filter. A 2.5 µL aliquot of the bacterial culture was added to each hydrogel of both stiffness conditions. The hydrogels were then polymerized followed by a 4 h growth at 37°C. A planktonic culture was started simultaneously as a control. To determine the colony-forming units (CFU mL⁻¹) before antibiotic treatment, hydrogels were mechanically homogenized to remove the cells, which were then plated on LB agar plates. Another set of hydrogels was treated with antibiotics, including colistin for 2 h, tobramycin for 2 h, meropenem for 5 h, and ciprofloxacin for 1.5 h. Following treatment, single cells were extracted by breaking hydrogels which were subsequently plated on LB agar plates. The number of colonies formed was used to determine the CFU mL⁻¹ values. Bacterial survival was assessed using log reduction plotting to quantify how effectively antibiotic treatment reduced bacterial populations. A higher log reduction value indicates greater bacterial killing, with each 1 log reduction representing a 10-fold decrease in bacterial count. The log reduction was calculated using the formula:

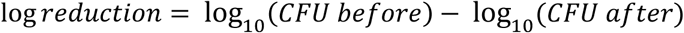

Planktonic cultures were also processed and analyzed using the same method.

#### **(ii)** Fluorescent imaging with Live/Dead staining

Planktonic cultures of the PAO1 WT strain labeled with *mNeonGreen* were embedded in hydrogels and incubated under the same conditions as described in section (i). After 4 hours of growth, antibiotics were added to the hydrogels. Colistin treatment was monitored at 2 and 4 h, while tobramycin, meropenem, and ciprofloxacin treatments were carried out for 20 h. Thirty minutes before imaging, propidium iodide (10 µM) was added and incubated to stain dead cells. Fluorescence imaging of antibiotic-treated planktonic cells (as shown in Figures 1 and 2) was performed using a 40X objective, capturing images at 2 h and 4 h for colistin, and at 20 hours for tobramycin, meropenem, and ciprofloxacin.

To visualize single cells, 1 µL of culture containing *mNeonGreen*-labelled cells and propidium iodide was carefully placed on a 0.8% agarose pad within a MaTek dish. Z-stacks were captured at 3 μm step size, covering a total depth of 30 μm to visualize the distribution across different z-planes. Live cells appeared green, and propidium iodide-stained dead cells are in magenta.

Image analysis was performed using Fiji. To quantify the percentage of dead biomass, the Otsu threshold was applied to the live/dead cell channels to accurately distinguish fluorescent signals. The areas of positive green pixels, representing live cells, and the propidium iodide positive pixels, indicating dead cells were measured. The percentage of dead biomass was calculated as the ratio of dead biomass to total biomass.

High-resolution images of planktonic and 3D bacterial clusters were captured using a 100X oil-immersion objective using the confocal spinning disk microscope. Imaging was performed at 2 h for colistin treatment and at 5 h for tobramycin, meropenem, and ciprofloxacin.

### Confocal imaging of 3D bacterial clusters and quantification of 3D Volume

PAO1 WT strain with constitutive *mNeonGreen* expression was grown to exponential phase (OD 0.5-0.6) in LB-medium from an overnight culture. The culture was then filtered and then embedded in PEG hydrogels of varying stiffness (1kPa, 11 kPa, 23 kPa and 63 kPa). Once confined, the cells grew into clusters over time, which were imaged using a 40X air objective on a Nikon Ti-2 spinning disc confocal microscope. For volumetric analysis, z-stack imaging with a 1 μm step size was set, capturing a total depth of 15 μm from the top of the hydrogel. The LB medium was refreshed at 3, 6, and 10 h, and imaging was performed across multiple fields of view and biological replicates at 3, 6, 10 and 20 h respectively. To quantify 3D biofilm volume, we employed a custom-developed Python script to pre-process z-stacks of 3D bacterial clusters. Before running the script, background subtraction was performed in Fiji using a 50-pixel rolling ball radius to remove background noise. The custom script applies a 3D median filter (5×5 in XY) to further reduce noise, followed by Li’s thresholding method with pixel values within the range of 70–65535. Connected component analysis with 26 connectivity was then used to label distinct clusters in 3D, retaining only those spanning at least four consecutive slices. To refine segmentation, a distance transform algorithm was computed, followed by watershed segmentation to accurately separate cluster boundaries. Finally, objects touching the image edges were cleared, and the processed images were saved as TIFF files. The processed TIFF files were then imported into the 3D Volume plugin within the 3D Suite in Fiji to quantify embedded biofilm volume. Volumetric measurements of biofilms within hydrogels of varying stiffness were used to generate frequency distribution plots and track volumetric cluster growth over time.

A similar approach was applied to PAO1 wildtype and Tn-seq mutants specifically Δ*ssg* and Δ*shaC* (PA1056), for imaging and quantification. Quantification was done as described above, except imaging was performed at a depth of 30 μm and the growth time was 4 h under confinement.

### Cell-length analysis post confinement

PAO1 WT clusters were grown for 6 h within PEG hydrogels (see Hydrogel Assay section for details). Single cells were mechanically extracted using a microblade and pestle in an Eppendorf tube, then resuspended in 1X PBS buffer. A 1 μL aliquot of the bacterial suspension was placed onto 0.8% PBS-agarose pads on Matek dishes. Phase-contrast images of individual cells were acquired using a Nikon Ti-Eclipse widefield microscope equipped with a 40X/0.75 Ph2 air objective. Cell length measurements were performed using BacStalk software. Before segmentation, images were enhanced through bandpass filtering. Individual cells were identified and segmented based on their boundaries, using a minimum threshold of 6 pixels and a maximum of 11 pixels. BacStalk automatically detected the medial axis of each cell, assigned a unique cell ID, and quantified cell length (in μm) as the distance along this axis. Manual corrections were applied when necessary to ensure measurement accuracy. The resulting cell length data was exported and analyzed using a custom Python script to generate plots, displaying individual measurements in colored points and black line marking the medians.

### High-resolution imaging of FM 4-64 stained 3D bacterial clusters

Wildtype *P. aeruginosa* cells were confined in PEG hydrogels of varying stiffness, as detailed in previous sections. After 6h, bacterial clusters were stained with FM4-64 dye (in the ratio 1:33) (N-(3-Triethylammoniumpropyl)-4-(6-(4-(Diethylamino) Phenyl) Hexatrienyl) Pyridinium Dibromide) for 15 minutes. High-resolution, single-plane images were taken using a fluorescence confocal scanning microscope. Post-processing was performed using the built-in AI feature to minimize noise, enhancing visualization of the cell membrane. Using a custom script integrated with Omnipose, we performed preliminary tests for single-cell segmentation within bacterial clusters and quantified bacterial cell lengths.

### Cell division and cell death in bacterial clusters

PAO1 WT cells were grown to the exponential phase (OD600 0.5-0.6) and embedded within PEG hydrogels of stiffness: 1kPa, 11 kPa, 23 kPa and 63 kPa. Hydrogels with bacteria were incubated at 37 °C to form clusters for 6 h. Single cells were recovered by breaking hydrogels mechanically using a pestle. The recovered cells were then plated on LB agar, and colony-forming units (CFU/mL) were measured to estimate the number of cell divisions under confinement. Colony forming units/mL were noted down the next day by counting the number of colonies. A planktonic culture was also incubated under the same conditions for 6 h as a control. Additionally, the initial cell inoculum (cells added to hydrogels at time = 0 h) was also accounted for in CFU measurements.

For cell death measurements, *P. aeruginosa* was grown under confinement for 6 h in hydrogels (1 kPa, 11 kPa, 23 kPa and 63 kPa) followed by propidium iodide staining for 15 min. The hydrogels were washed with 1x PBS solution to remove excess dye. Z-stacks of hydrogels were captured using the 40x air objective on a Nikon Ti2-eclipse spinning disk confocal microscope in both green and red fluorescence channels. The percentage of dead biomass was quantified by measuring positive green pixels (indicating live cells) and positive red pixels (highlighting dead cells). The fraction of dead biomass was then calculated using the formula:

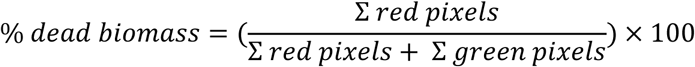

### Growth experiments of confined cells

The exponential culture of PAO1 WT cells was embedded in 1 kPa and 23 kPa hydrogels and were grown for 4 h at 37°C. Next, the gels were manually homogenized using a pestle to recover the cells. 1 µL aliquot of the recovered cells was placed on a 0.8% LB agarose pad, and phase-contrast time-lapse images were recorded for 2 h. Imaging was performed using a 100X oil objective Nikon Ti2-Eclipse spinning disk microscope, equipped with an incubation chamber adapter to maintain the temperature at 37°C. Phase-contrast movies were analyzed using Omnipose to segment individual cells in the movie. The segmented movies were then imported on Trackmate cell tracking. Using the overlap tracker option (with IoU calculation set to ‘precise’, a minimum IoU value of 0.1 and Scale factor of 1.3) were able to get tracks of individual cells growing over time. The area of individual cells was measured and exported as CSV files. To quantify the elongation rate, a custom Python script was used. Filtering criteria were applied to ensure that only single cells present in frame 0 were tracked until their first division. Additionally, only cells that were segmented for more than two frames were included in the analysis. The normalized area of each cell was plotted over time. The growth rate (*r*) is calculated using the equation

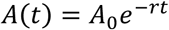

Where *A(t)* is the area of the cell at time *t, A0* is the initial area of the cell, and *r* is the growth rate. Exponential fit on each individual tracks, the slope *r* was extracted, which represents the growth rate of cells over time.

### Single-cell kinetics assay for antibiotic-induced killing

Planktonic PAO1 WT cultures were embedded in hydrogels, incubated, and removed from hydrogels as described in the previous section (regrowth experiments). A 1 μL sample of recovered cells was placed on a 0.8% LB agarose pad containing antibiotics and immediately imaged using a 40X air objective on a Nikon Ti2-E Eclipse spinning disk confocal microscope. For colistin, images were captured every 5 min over 2 h using the 488 nm laser to capture the green fluorescence channel. For ciprofloxacin, images were taken every 15 min for a total duration of 10 h. The survival population was quantified by counting the number of cells at each time point, and the killing percentage was determined by normalizing the number of cells to the initial time frame.

### Transposon sequencing experimental design

We performed two distinct Tn-seq experiments: **(A)** a screen searching for P. aeruginosa genes for growth under mechanical confinement, and **(B)** a screen looking for genes involved in antibiotic tolerance under these conditions.

#### **(A)** Tn-seq for constrained growth in PEG hydrogels

We used a previously published *P. aeruginosa* PAO1 transposon mutant library generated by conjugating the Tn5-based transposon T8 (IS*lac*Z*hah*-tc) into the PAO1 genome, as described in references^62^. The library consists of approximately 250,000 unique mutants and was stored in LB with 15% glycerol at -80 °C at an OD600 of ∼56. The library stock was diluted in LB to an OD600 of 0.2 (1.5 mL) and grown until the exponential phase (OD600 -0.6) for 2 h. Then, the cells were spun down, washed in 1x M9 buffer, resuspended in 150 μL of PBS and used as inoculum for embedding them in PEG hydrogels. To ensure sufficient cell number for sequencing, keeping a high density of inoculum was important. The mutant library was embedded in PEG hydrogels with stiffness 1 kPa and 23 kPa, and 20 hydrogels (10 Matek dishes used) were prepared per condition to have approximately ∼ 10^6^ *P. aeruginosa* cells in total. Growth under confined conditions was carried out in 3 mL LB medium at 37 °C in each dish. After 4 h of growth, all 20 hydrogels in each stiffness condition were pooled together and homogenized in 300 μL of LB medium supplemented with 0.1 mm size glass beads using a homogenizer. Homogenization was performed for a total of 1 minute in 15-sec bursts, with 5- sec colling intervals between each burst. Following this step, samples were re-grown in 1 mL LB medium for an additional 2 hours at 37 °C with constant shaking. Cells were then separated from hydrogel debris by passing the suspension through a 5 μm filter, followed by centrifugation at 14000 rpm for 2 min. The resulting pellets were washed twice in PBS to remove LB traces, and stored at -80 °C until subsequent processing.

In parallel, we prepared planktonic LB control samples using the same inoculum. One control was exposed to UV light for 1 min to mimic the conditions used during hydrogel polymerization, while the other was grown without UV exposure in 3 mL LB at 37 °C. We grew both controls for 4 h in parallel with the confined samples. After this initial growth, we centrifuged the 3 mL cultures, resuspended in 1 mL of fresh LB, and continued for an additional 2 h. Finally, cells were spun down, washed in PBS twice, and the pellet was stored at -80 °C until further processing.

#### **(B)** Tn-seq for antibiotic tolerance under mechanical confinement

Growth of the Tn-seq mutant library in PEG hydrogels was set identically to the Tn-seq above, with the following 3 experimental conditions: (1) untreated planktonic cells (negative control), (2) planktonic cells treated with the antibiotic, and (3) cells embedded in 23 kPa hydrogel. All conditions were incubated at 37 °C for 4 h. For antibiotic exposure in 23 kPa hydrogel condition, colistin was added directly to the dish at a concentration of 5 μg mL^-1,^ and exposure lasted for 60 min. For tobramycin treatment, we used a final concentration of 10 μg mL^-1,^ with a 20-min exposure. These concentrations and exposure times were defined after the MIC assay and were chosen to induce approximately two logs of killing in embedded clusters, as established in prior tolerance assays. Following antibiotic treatment, hydrogels were homogenized in 300 μL of fresh LB medium as described above. An outgrowth step was then performed in 1 mL of LB: colistin-treated samples were incubated for 3 h, while tobramycin-treated samples required 11-12 h to reach an OD600 of 1.5-1.9. samples were subsequently passed through a 5 μm filter to separate cells from hydrogel debris, and cells were pelleted by centrifugation at 14000 rpm for 2 min. The same protocol was applied to untreated and antibiotic-treated planktonic cells, except for the homogenization step. To match the final OD600 of hydrogel samples, regrowth of planktonic samples was limited to 2 h before harvest.

### Tn-Sequencing

Genomic DNA from *P. aeruginosa* samples (185 ng) was first sheared with a Covaris S220 using 400 bp insert settings (50 uL in microTUBES with AFA fiber, Peak incident power: 175, Duty factor: 5%, Cycles per burst: 200, time: 55 s). Libraries were prepared with the xGen DNA MC Library Prep Kit (IDT, protocol version v2) using xGen UDI-UMI adapters (IDT, 15 µM stock). With these adapters, P5 and P7 sequence are inverted compared to Illumina adapters, allowing transposon sequencing directly from read 1 (P5 side) in a single-end run (see below). The purified ligated product was PCR amplified with a primer specific for the Illumina P7 sequence (CAAGCAGAAGACGGCATACGA) and a second one specific for the transposon sequence (cgacgttgtaaaacgaccacgt) carrying a 5’-biotin. PCR was performed with the KAPA HiFi HotStart ReadyMix kit (Roche). Cycling conditions were 95°C for 3 min, followed by 12 cycles of 98°C for 15 s, 60°C for 30 s, and 72°C for 30 s, and a final extension of 1 min at 72°C. The library was purified with SPRI beads at a 1X ratio. The PCR product was captured with pre-washed Dynabeads MyOne Streptavidin T1 (ThermoFischer). Binding & Wash (B&W) Buffer 2X composition is 10mM Tris-HCl pH 7.5, 1mM EDTA, 2M NaCl. At least 1ml of 1X B&W Buffer was added and mixed with 25 µl of Dynabeads. After 1min on a magnet, the supernatant was removed and Dynabeads washed with the same volume of 1X B&W Buffer. This wash was repeated a second time. After removal of the supernatant on the magnet, the Dynabeads were resuspended with 50 µl of 2X B&W Buffer.

50 µl of library was mixed with the washed Dynabeads followed by incubation at RT on a rotator for 30 min. After 2 min on a magnet and discard of the supernatant, a wash was done with 100 µl of 1X B&W Buffer. Two additional washes were done before the final elution in 40 µl H2O. Half of the washed capture was used for the nested PCR with the Illumina P7 sequence (See sequence) and a tailed primer made of the Illumina P5 sequence (underlined), the TruSeq read 1 primer binding site (bold), and a transposon specific binding sequence <u>AATGATACGGCGACCACCGA</u>**GATCTACACTCTTTCCCTACACGACGCTCTTCCG**ATCT*c caggacgctacttgtgtat*. This nested PCR amplification was performed with the KAPA HiFi HotStart ReadyMix kit with same cycling conditions as above but 10 cycles. The final library was purified with SPRI beads at a 0.7X ratio. It was quantified with a fluorometric method (QubIT, Thermo Scientifics) and its size pattern analyzed with a fragment analyzer (Agilent). Sequencing was performed on a Aviti (Element Biosciences) on a Cloudbreak Freestyle high output flow cell for a 150 cycles single end sequencing run. Clustering was performed with 1nM library spiked with PhiX (Element Biosciences). Base calling and demultiplexing was done with bases2fastq version: 2.0.

### Tn-seq analysis

For each sample, we first merged demultiplexed raw reads from two sequencing lanes into a single file. The resulting reads were then processed and aligned to the *Pseudomonas aeruginosa* PAO1 genome (NCBI: NC_002516) using the TPP pipeline, available in TRANSIT v3.2.7, with BWA v0.7.17 as the aligner. Mapping was performed using the ‘mem’ algorithm and the ‘Tn5’ protocol, along with default TPP parameters (e.g., ‘Max reads’ set to −1, ‘Mismatches allowed’ = 1, and a prefix search window of 0, 20)^86,87^ .Following alignment, we used TRANSIT to conduct quality control and downstream analysis of the generated ‘wig’ files. Because the dataset displayed notable skewness, we applied the Beta-Geometric Correction (BGC) for normalization, using the ‘normalize’ command with the ‘-n aBGC’ flag as recommended in the software documentation. To evaluate conditional gene essentiality across experimental conditions, we applied the ‘resampling’ method within TRANSIT (run in GUI mode), specifying the Tn5 transposon and retaining most default settings (10,000 samples, pseudocount = 0.00, adaptive resampling = True, histogram = False, include Zeros = True). Since normalization had already been applied, no additional normalization was selected during the resampling step. We assessed the following condition pairs, with the first condition in each set treated as the experimental group and the second as the control: (1) LB control (no UV) vs. LB (UV treated), (2) stiff gel vs. LB control (UV treated), (3) stiff gel (colistin-treated) vs. stiff gel untreated, (4) stiff gel (tobramycin treated) vs. stiff untreated, (5) liquid treated (colistin-treated) vs liquid untreated, and (6) liquid treated (tobramycin treated) vs liquid untreated. Comparisons 1–2 correspond to the growth under confinement Tn-seq dataset, while comparisons 3-6 relate to the antibiotic tolerance in stiff gel conditions Tn-seq. While sequencing replicates were not generated for each sample, the input for each condition consisted of pooled material from ten hydrogels, helping to capture biological variability. Statistical significance was assessed using an adjusted *p*-value threshold of <0.05 (Benjamini– Hochberg correction for false discovery rate), in accordance with TRANSIT recommendations. As an additional control, we repeated the resampling analysis using trimmed total reads (TTR) for normalization. Although TTR normalization also identified many of the same significant genes, it tended to yield a higher number of hits, including genes with low log₂ fold-change values. Given the known advantage of BGC in minimizing false positives from skewed Tn-seq data, we relied primarily on the more conservative BGC-normalized results. Due to the limited propagation potential of our library (constrained by material/inoculum size), we chose not to apply a log₂ fold-change cut-off. All genes meeting the adjusted *P* < 0.1 (growth) and *P* < 0.05 (antibiotic-related comparisons) threshold are reported.

### Growth and Antibiotic tolerance assay of Tn-seq mutants

PAO1 wildtype and mutant strains (Δ*amgS*, Δ*phoQ*, Δ*shaC*, and Δ*ssg*) were grown to the exponential phase (OD600 0.5–0.6) and embedded in 23 kPa PEG hydrogels. Hydrogels containing bacteria were incubated at 37 °C for 4 h to allow cluster formation. In parallel, planktonic cultures of both wildtype and mutant strains were grown under identical conditions. After incubation, hydrogels were mechanically broken with a pestle to recover single cells. Recovered cells were plated on LB agar, and CFUs/mL were counted the following day to estimate cell divisions under confinement. The initial inoculum was also plated to account for starting CFU levels. Growth (generations in 4 h) was quantified as the log2 fold change in CFUs between the initial inoculum and final number of cells at time = 4 h.

For tolerance assay, bacterial clusters were grown for 4 h before antibiotic exposure. Colistin- related mutants were treated with colistin for 1 h. For tobramycin treatment, clusters were exposed for 4 h for imaging experiments and 30 min for CFU-based assays. For fluorescence imaging, Syto 9 and propidium iodide (PI) were added to the samples after washing away the antibiotics. PI was used at a final concentration of 5 μM, while the concentration of Syto 9 was determined separately. Z-stacks were acquired to a depth of 50 μm with a step size of 3 μm. Multiple regions of interest (ROIs) were imaged across biological replicates. Dead biomass was quantified by measuring green (Syto 9–positive) and red (PI-positive) pixels. Percentage cell death was calculated as the ratio of PI-positive pixels to the total pixel count (Syto 9 + PI).

### Quantification of deformation pressure in growing bacterial clusters

Bacterial clusters embedded in PEG hydrogels were treated as expanding spherical cavities within an hyperelastic solid to estimate the mechanical pressures generated during growth. The cavitation model from Barney *et al*^55^ describes the cavity pressure as equation (1):

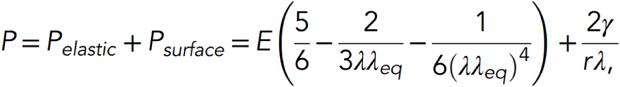

Here, *P* is the internal cavity pressure, *E* is the hydrogel’s elastic modulus, λ = r/r0 is the expansion ratio, λeq accounts for the balance between interfacial energy and elasticity, γ is the interfacial energy, and *r* is the undeformed cavity radius. For analysis, surface energy effects were not included, setting *Psurface* = 0, which makes γ = 0 (no surface energy) and λeq = 1. This simplified, and the equation (2) is:

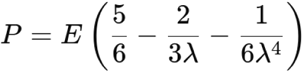

Cluster volumes were measured experimentally at 3, 6, 10, and 20 hours. The initial cluster volume was set to 3 μm^3^. Assuming spherical geometry, we converted these volumes into radii and lambda (λ) was calculated. This experimentally determined λ was then used in in the simplified cavitation model (Equation 2) to calculate the internal pressure generated by bacterial clusters at each time point, using the known elastic modulus *E* of the PEG hydrogel.

### Data analysis and Statistical analysis

Most statistical analyses and plots were done with custom Python scripts. Image analyses were mainly carried out in Fiji. Some additional statistical tests, plots, and image analyses were done using GraphPad Prism (version 8) and Python.

## Authors contributions

Conceptualization: SM, SS, AP Data curation: SM

Formal Analysis: SM Funding acquisition: SS, AP Investigation: SM Methodology: SM, ZM Project administration: AP Supervision: SS, AP Visualization: SM

Writing – original draft: SM, AP

Writing – review & editing: SM, SS, AP

## Acknowledgements

We thank Alice Cont and Florent Prongué for their assistance with modulus measurements, and Prof. Ester Amstead for providing access to the rheometer. We thank Dea Müller for designing primers for the deletion mutants. We acknowledge the Lausanne Genomics Technology Facility (GTF) for assistance with Tn-seq sequencing and initial data processing. We are grateful to Lucas Meirelles for discussions on Tn-seq experiments and to Laure le Blanc for sharing the python script for single-cell tracking analysis. We also thank Prof. Nikolas Bouklas (Cornell University) for providing the Python script for pressure calculations.

