## Supplementary material for "Growth in confinement promotes *Pseudomonas aeruginosa* tolerance to antibiotics": Figure S1-15, Table S1-5

#### Supplementary figures:

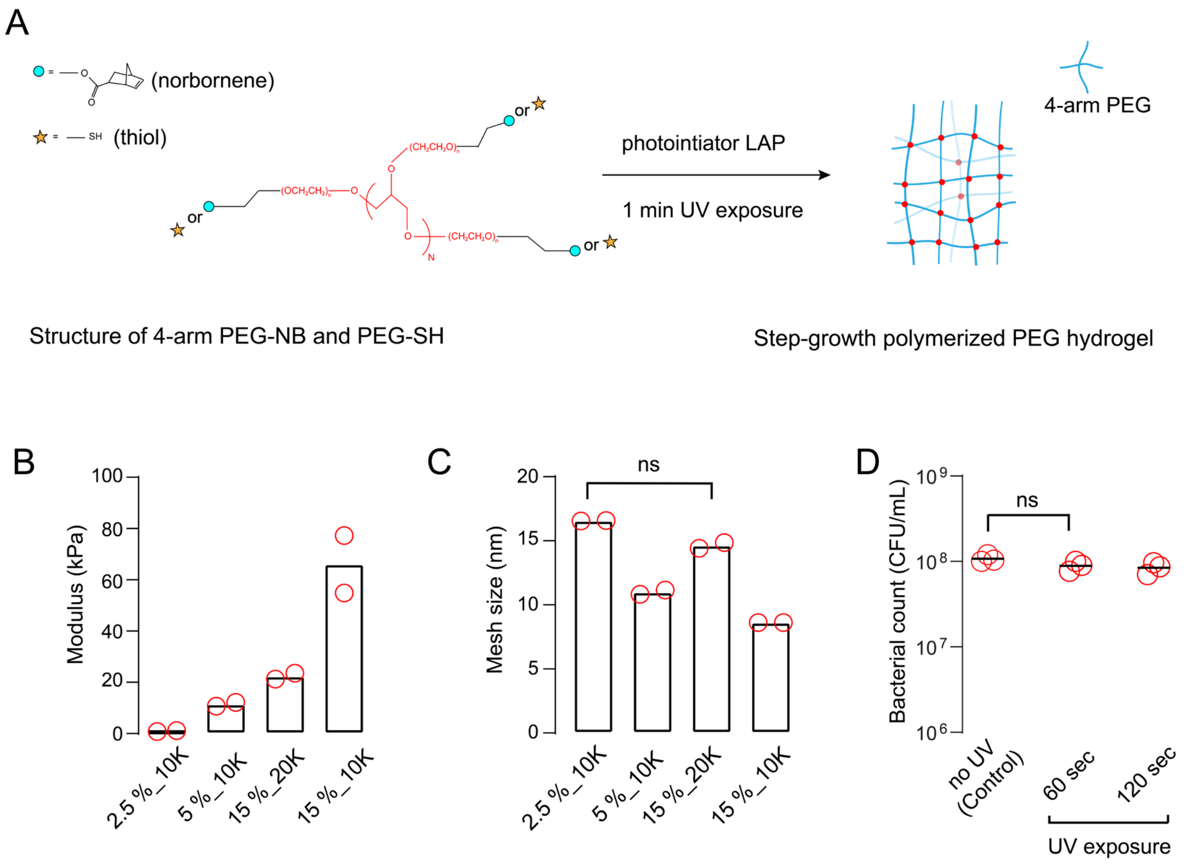


**Figure S1 Mechanical characterization of PEG hydrogels**. **(A)** Chemical structures of PEG-norbornene and PEG-thiol monomers, and the schematic of photo crosslinking step forming the hydrogel network. The blue lines represent the 4-arm PEG monomers, while red points indicate the crosslinks formed upon UV polymerization in the presence of the LAP photoinitiator. **(B)** Young's Modulus measurements showing the effects of varying PEG molecular weight (10 and 20 kDa) and concentration (2.5 %, 5% and 15%) on hydrogel stiffness. Each red circle indicates a biological replicate (N=2), with each comprising 3 technical replicates. **(C)** Mesh size measurements of PEG hydrogels with different MW and concentration, N=2 ($p$=0.333, Mann-Whitney test). **(D)** Survival of bacterial population upon UV light exposure. Data from three biological replicates (N=3), each with three technical replicates ($p$ = 0.1088, Welch's $t$-test).

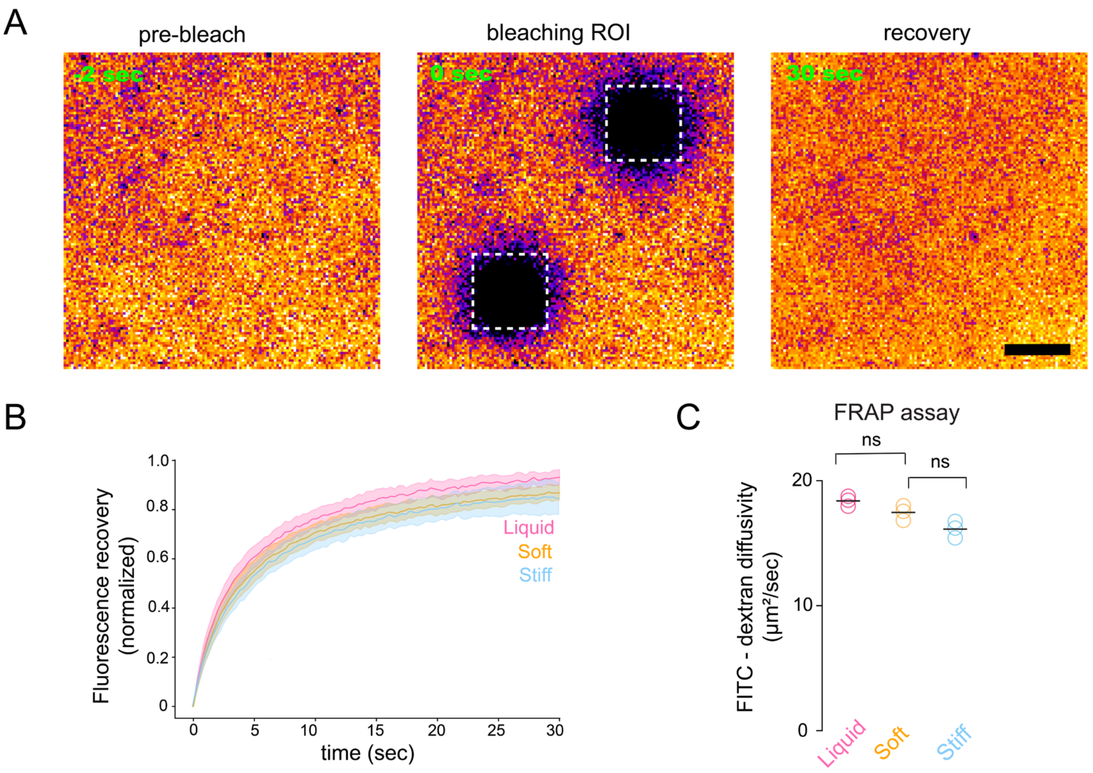


**Figure S2 FRAP assay for diffusion. (A)** Confocal microscopy images of FITC-dextran-loaded PEG hydrogel showing pre-bleach, post-bleach (0 s) and recovery (30 s) frames. Scale bar 100 μm. **(B)** Recovery curves comparing liquid, soft and stiff hydrogels. For all the curves, time zero ($t$ = 0 s) marks the post-bleach frame where recovery begins. **(C)** FRAP-based diffusion measurements of FITC–dextran (10 kDa) in liquid and PEG hydrogels, including both soft (1 kPa) and stiff (23 kPa). Diffusion rates quantified from recovery curves, showed no significant differences across all conditions. Data represent 3 biological replicates with 3 technical replicates each. Statistical analysis performed using unpaired Welch's $t$-test, $p$ = 0.087 (liquid vs 1 kPa), $p$ = 0.059 (1 kPa vs 23 kPa).

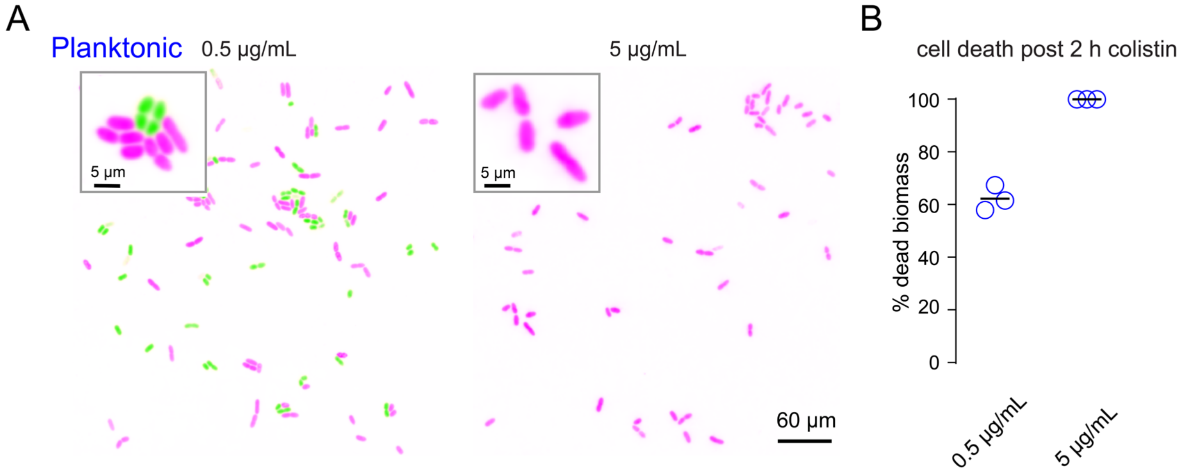


**Figure S3 Cell death upon colistin exposure**. (**A**) Confocal images of planktonic *P. aeruginosa* after colistin-treatment under at concentrations, 0.5 μg/mL, and 5 μg/mL. Live cells are in green, while dead cells in magenta (Scale bar: 60 μm). Inset shows cell death at single-cell level (Scale bar: 5 μm). **(B)** Dead biomass quantification of planktonic cells exposed to two different colistin concentrations (number of bacteria > 100 from each biological replicates, N= 3).

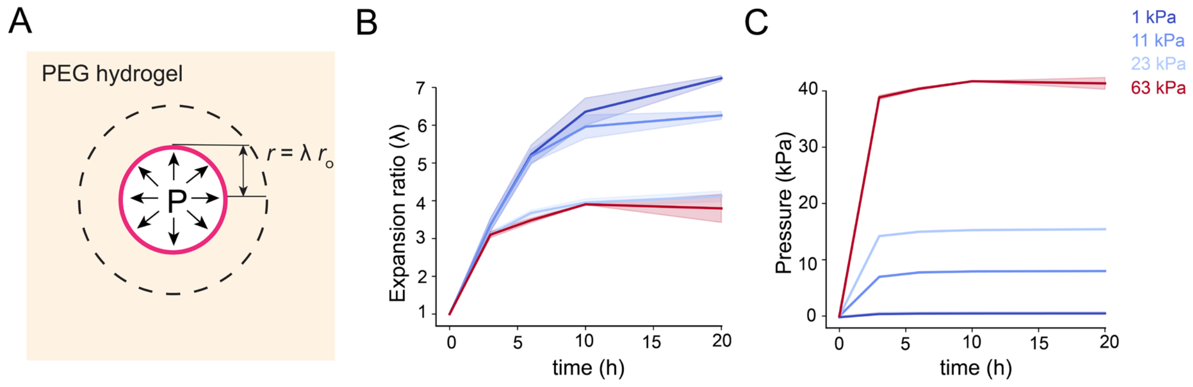


**Figure S4 Cavitation pressure estimation. (A)** Schematic showing spherical cavity representing a bacterium in PEG hydrogel that experiences an increase from initial radius $r$ to final radius $r_0$. P is the growth-induced pressure generated by bacteria and λ is expansion ratio ($r_0$/r) calculated using before and after volumetric growth **(B)** Expansion ratio (λ) measurements of bacterial clusters over time. **(C)** Pressure calculations (kPa) of growing cavity at different time points (h). Color code represents the Young's Modulus of the PEG hydrogels used in the experiments**.**

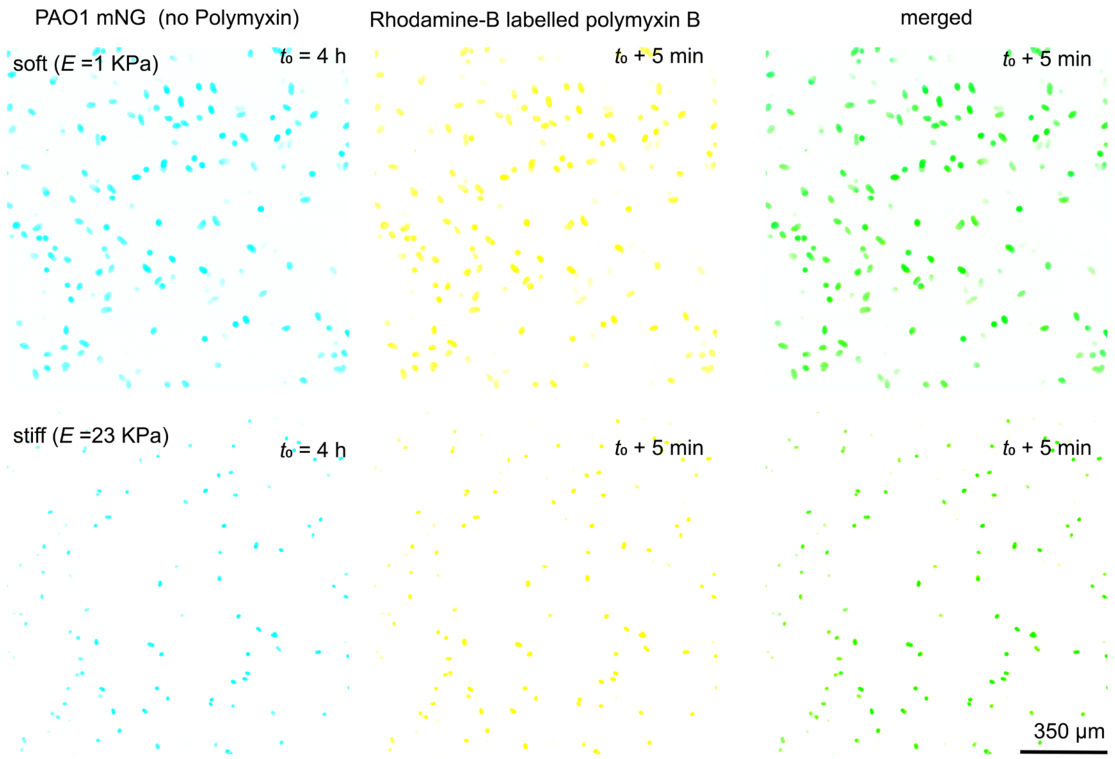


**Figure S5 Antibiotic uptake assay.** Fluorescence images of PAO1 bacterial clusters grown for 4 h in soft and stiff hydrogels (left). Visualization of antibiotic uptake at $t$ = 5 min (middle section) using Rhodamine-B polymyxin, indicating the penetration of antibiotic through both hydrogel of different stiffness and reaching the clusters. Scale bar: 350 μm.

A

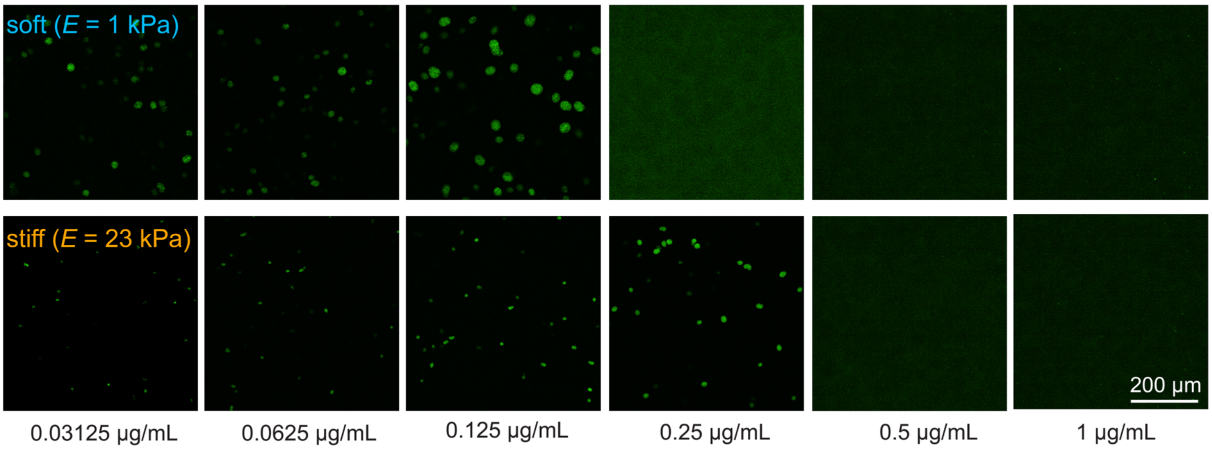


B

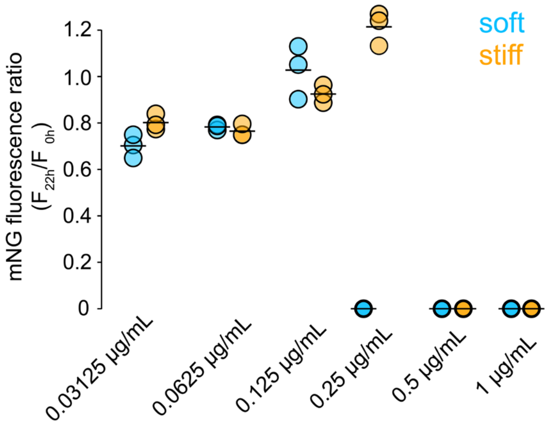


C

| | **Minimum inhibitory concentration (MIC) for colistin (μg/mL)** |
|---|---|
| Planktonic | 0.125 |
| soft (*E* = 1 kPa) | 0.25 |
| stiff (*E* = 23 kPa) | 0.5 |

**Figure S6 Embedded MIC (Minimum inhibition concentration) assay with colistin. (A)** Confocal fluorescence images of *PAO1::mNeonGreen* clusters grown in soft and stiff hydrogel treated with colistin from time 0 h. Green fluorescence shows bacterial growth after 22 h. Scale bar: 200 μm. **(B)** The ratio of *mNeonGreen* fluorescence intensity at 22 h /0 h was used as a viability metric. A ratio of 0 indicates no detectable survival, while ratios more than 0.5 indicate substantial bacterial survival at the tested colistin concentrations. **(C)** Table showing the minimum inhibitory concentration (MIC) measurements of PAO1 under planktonic and confined conditions when exposed to colistin.

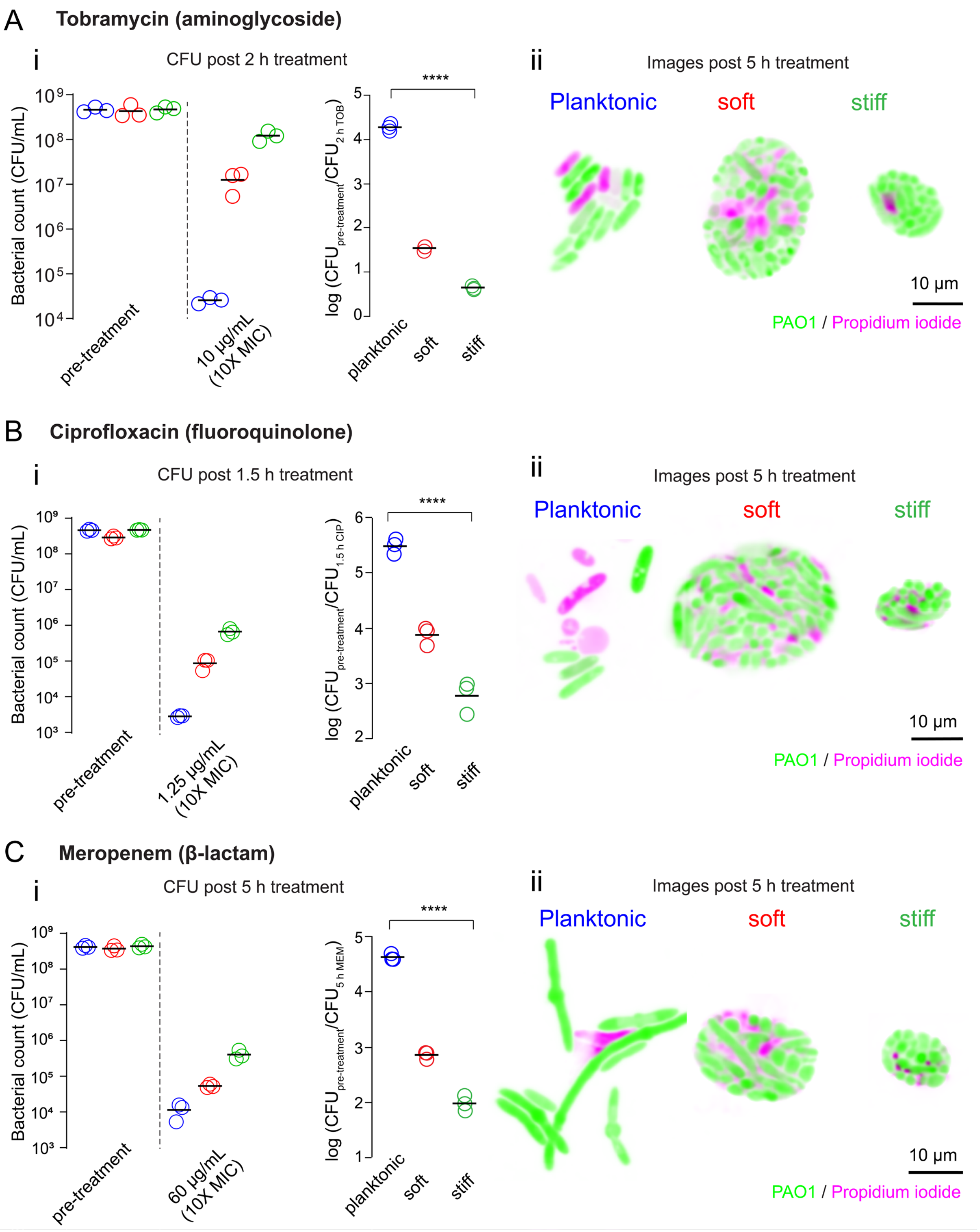


**Figure S7 Confined growth mechanically promotes tolerance across antibiotic classes.** PAO1 clusters grown for 4 h in hydrogels before treatment with 3 different antibiotic classes. **(A, i)** Tobramycin treatment (10 µg/mL) for 2 hours. Absolute survival determined using CFU counts, with log-reduction plots quantifying the fold-change **(B, i)** CFU counts before and after 1.5 hours of ciprofloxacin (1.25 µg/mL) treatment on *P. aeruginosa* clusters under planktonic and confined conditions. **(C, i)** Impact of 5 h meropenem (60 µg/mL) treatment on cluster viability comparing CFU counts, and log-reduction highlighting the killing effect in all conditions. Statistical analysis of log-reduction between planktonic and stiff hydrogel conditions showed a significant difference for all antibiotic classes, with Tukey's HSD indicating $p < 0.0001$. For all the above-chosen antibiotics **(A-C, ii)**, representative 100X fluorescence images of antibiotic-treated PAO1 clusters after 5 h of exposure. Live cells are indicated in green, while dead cells are in magenta. Scale bar is 10 µm.

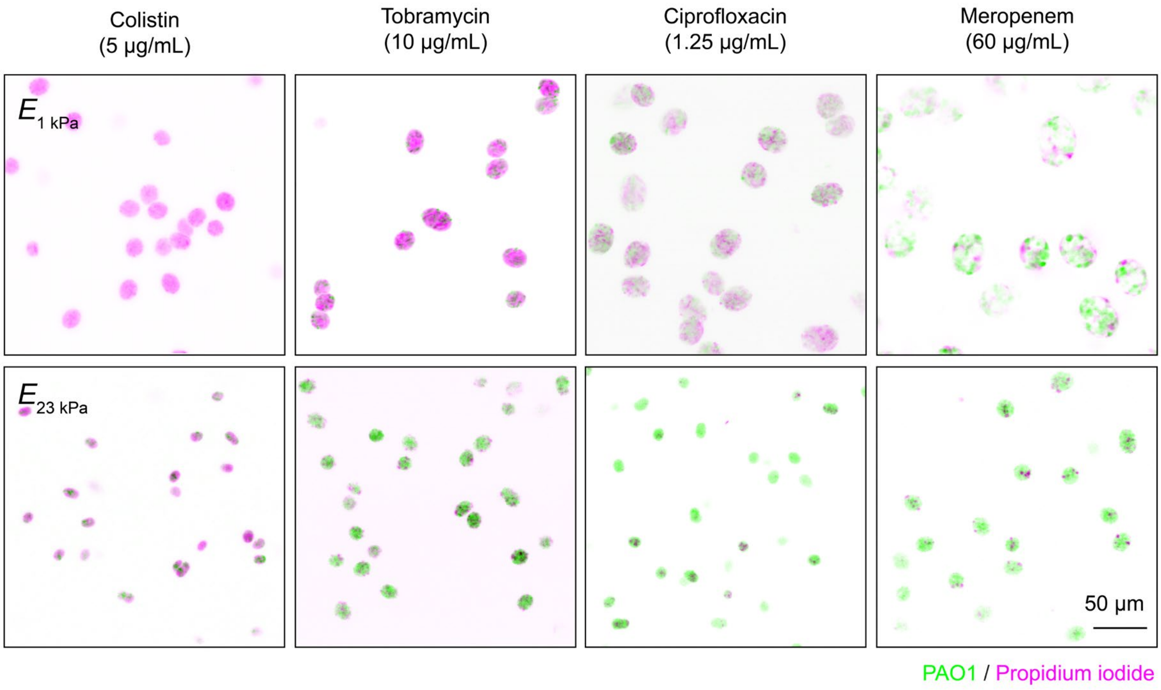


**Figure S8 Long-term effects of antibiotic exposure.** Fluorescent confocal spinning disk images of PAO1 clusters treated with clinically relevant antibiotics. Clusters were imaged after 4 hours of colistin treatment and 20 h post-treatment with tobramycin, ciprofloxacin, and meropenem. Scale bar: 50 µm.

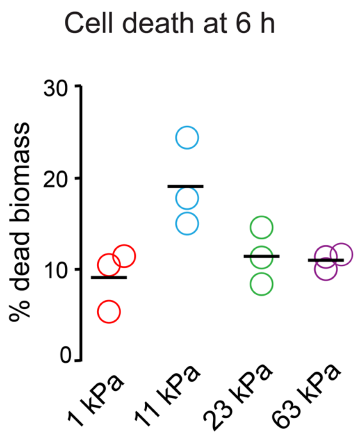


**Figure S9 Cell death in bacterial clusters**. Quantification of cell death in PAO1 3D clusters after 6 hours growth in hydrogels of varying stiffness, labelling dead biomass using propidium iodide stain. Each circle represents biological replicates (N=3) and black dash represents the mean of the biological replicates.

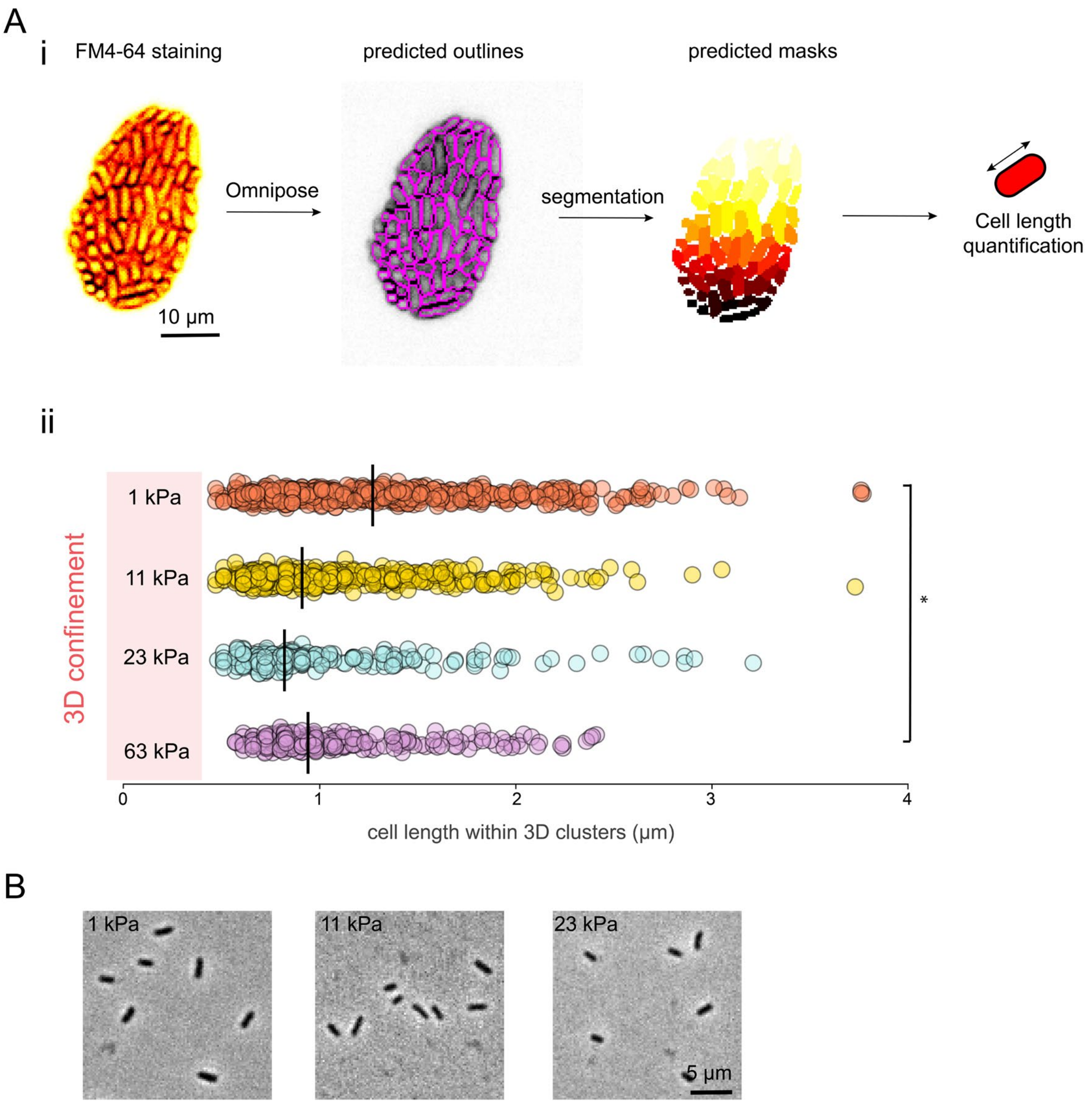


**Figure S10 Bacterial cell length quantification under confinement. (A, i)** Image analysis pipeline to quantify single-plane images of FM4-64-stained bacterial clusters inside hydrogels. This is done first by segmentation using Omnipose to obtain predicted outlines and masks, followed with quantification. **(A, ii)** Bacterial cell lengths quantification from the predicted masks using a custom Python script (Mann-Whitney t-test, **p* = 0.0127 between 1kPa vs 63 kPa stiffness). **(B)** Phase contrast images of single-cells removed from 1kPa, 11 kPa and 63 kPa hydrogels.

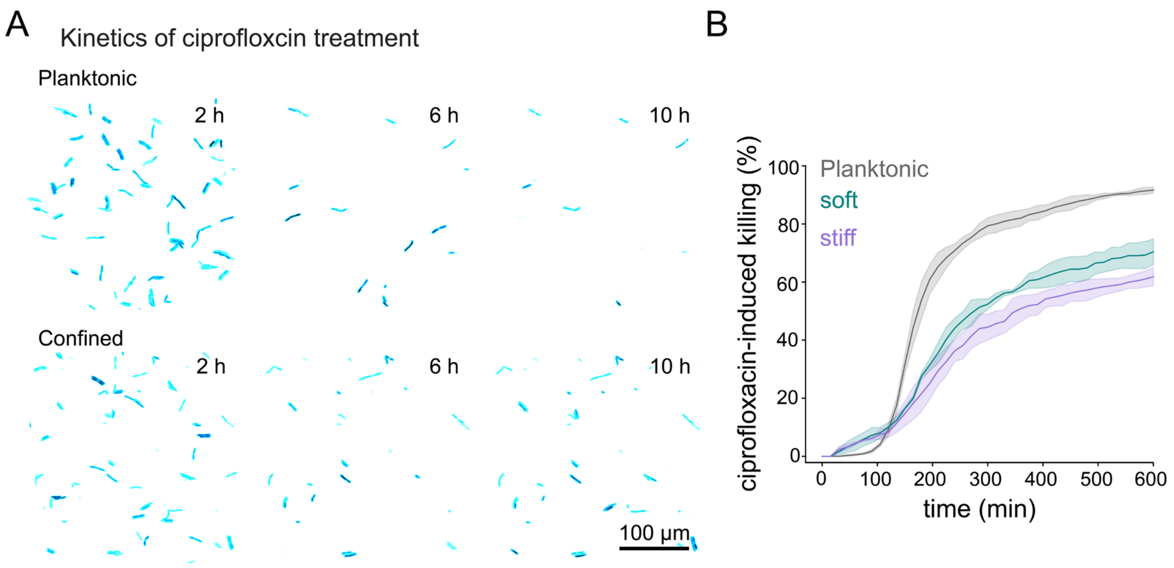


**Figure S11 Ciprofloxacin treatment of released cells after confinement. (A)** Time-lapse microscopy images comparing planktonic cells and confined cells from 1 kPa exposed to ciprofloxacin at at 2 h, 6 h, and 10 h. Scale bar = 100 µm. **(B)** Killing kinetics quantification for previously confined cells exposed to ciprofloxacin. Solid lines indicate the mean of the biological replicate means, and shaded areas show the standard deviation.

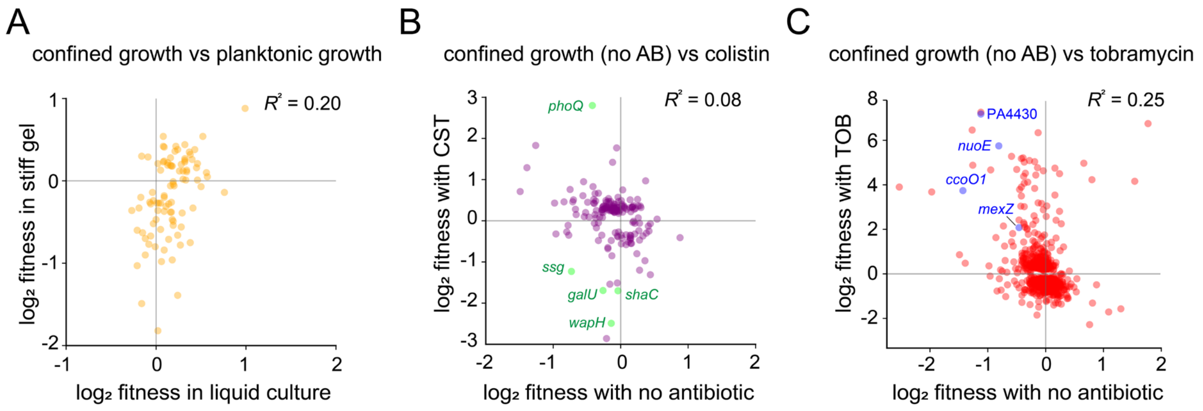


**Figure S12 Fitness correlations and gene ontology of confinement-specific determinants**. (**A**) Correlation between growth of *P. aeruginosa* mutants in gel and planktonic conditions**.** Each yellow data point represents a gene identified in the Tn-seq screen. Only genes that were significant in at least one condition (adjusted $p < 0.1$, resampling method in TRANSIT with Benjamini–Hochberg correction) are shown. $R^2$ indicates the coefficient of determination. (**B**) Fitness correlation of *P. aeruginosa* transposon insertions in colistin treated confined population versus untreated. Each data point represents a gene from the Tn-seq dataset under colistin treatment in confinement. Purple points show all genes, and green points highlight selected genes. (**C**) Scatter plots showing fitness correlation of *P. aeruginosa* transposon insertions under tobramycin confinement. Red dots show all genes; blue dots highlight specific genes under drug exposure. In the plots B and C, $R^2$ denotes the coefficient of determination.

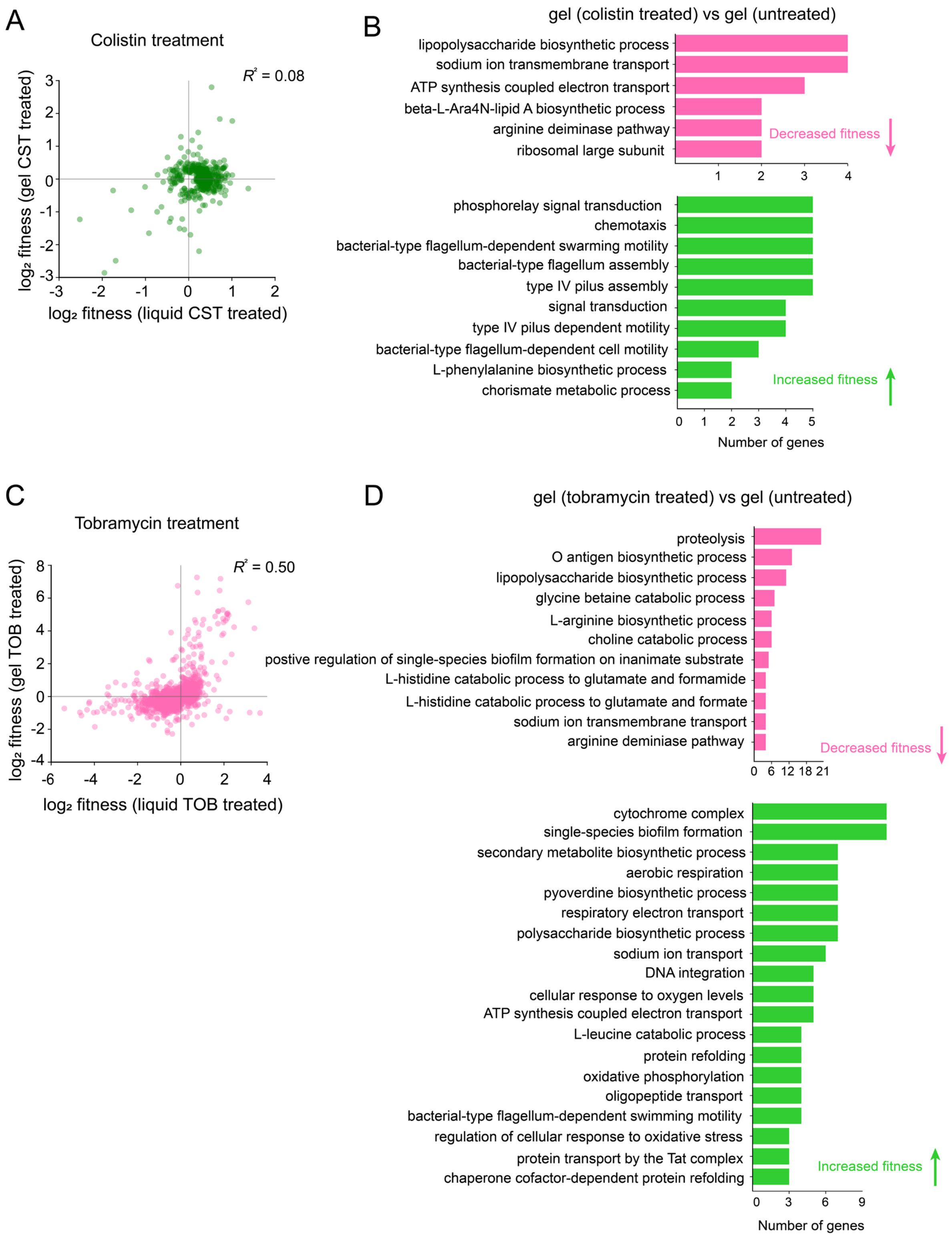


**Figure S13 Gene ontology analysis of antibiotic-treated bacterial clusters.** (**A**) Correlation between *P. aeruginosa* Tn-seq insertions in stiff gel and planktonic conditions under colistin exposure**.** Each green data point represents a gene identified in the Tn-seq screen. (**B**) Gene ontology analysis of biological processes using DAVID, comparing genes enriched or depleted in gel under colistin treatment relative to planktonic treated control. (**C**) Relationship between fitness of *P. aeruginosa* insertions in gel versus planktonic conditions during tobramycin treatment**.** Pink data points represent genes identified by Tn-seq. (**D**) DAVID gene ontology analysis of confined cells treated with tobramycin, compared planktonic treated controls. In both plots (**A**) and (**B**), only genes that were significant in at least one condition (adjusted $p < 0.1$, resampling method in TRANSIT with Benjamini–Hochberg correction) are shown. $R^2$ indicates the coefficient of determination. In **C** and **D,** top panel (pink) shows biological processes where mutants exhibit decreased fitness, while bottom panel (green) indicates increased fitness.

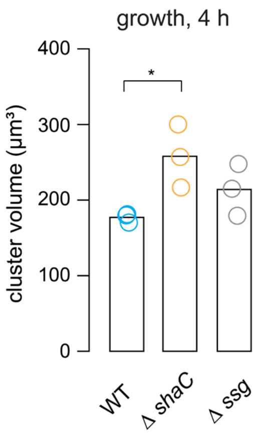


**Figure S14 Cluster volume quantification of Tn-seq mutants.** Bar graph showing the mean cluster volume after 4 h of growth in stiff PEG hydrogel. Scatter points indicate mean values (N=3 biological replicates). Tukey's MSD test show significance between wildtype (WT) and Δ*shaC*, **p* = 0.0449.

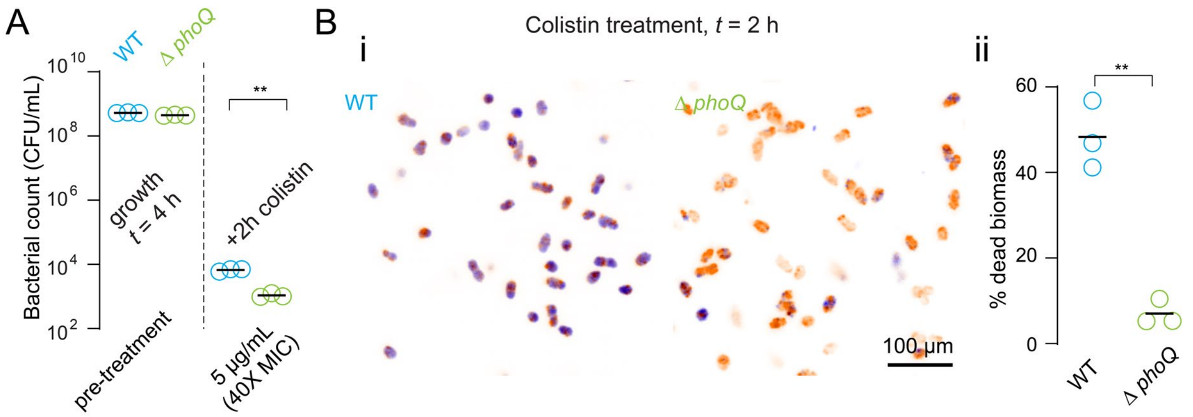


**Figure S15 Mutation in *phoQ* has opposite effect on tolerance in planktonic and confined populations. (A)** Survival after 2 h colistin exposure was measured by CFU counts for WT and Δ*phoQ* cells in liquid culture. Each point shows the mean of a biological replicate, and the black line marks the mean across replicates ($N$ = 3). Statistical comparison of wildtype and mutant after treatment, done with unpaired *t*-test with Welch's correction, ***p* = 0.0028). **(B, i)** Confocal maximum intensity projections of colistin-treated clusters. Images highlight comparison between wildtype and Δ*phoQ* mutant. Live cells are indicated in orange, while dead cells are in blue. Scale bar, 100 µm. (**B, ii**) Quantification of percentage dead biomass of clusters in gel to 5 µg/mL colistin (two-tailed non-parametric *t*-test with Welch's correction, ***p* = 0.0062).

### Supplementary tables

##### Table 1 Monomer and Crosslinker Ratios for PEG Hydrogel Preparation

| Concentration | PEG-SH (µL) | PEG-NB (µL) | M9 solution (µL) | Bacteria (µL) | LAP (µL) | Modulus (kPa) | Average water content (%) |
|---|---|---|---|---|---|---|---|
| 2.5% | 3.125 | 3.125 | 36.25 | 5 | 2.5 | 1 | 99.18 |
| 5% | 6.25 | 6.25 | 30 | 5 | 2.5 | 11 | 96.99 |
| 15% | 18.75 | 18.75 | 5 | 5 | 2.5 | 23 | 96.36 |
| 15% | 18.75 | 18.75 | 5 | 5 | 2.5 | 63 | 94.51 |

##### Table 2 List of bacterial strains and plasmids used in this study

| **Name** | **Description** | **Origin/reference** |
|---|---|---|
| PAO1 (ATCC 15692) | Persat lab wildtype strain (914) | |
| PAO1 *mNeonGreen* | PAO1 constitutively expressing mNeonGreen (attTn7::miniTn7T-Gm-ptet::mNeonGreen) | This study |
| PAO1 Δ*phoQ* | Deletion in *phoQ* | This study |
| PAO1 Δ*ssg* | Deletion in *ssg* | This study; strain in the lab |
| PAO1 Δ*PA1056* | Deletion in *PA1056* | This study |
| PAO1 Δ*amgS* | Deletion in *amgS* | This study |
| *E. coli* XL10-gold | Used for routine cloning and plasmid stocks | |
| pEX18Gm | Exchange vector for S17 mating | |

##### Table 3 List of primers

| **Primers** | **5'-3' sequence** |
|---|---|
| oSKJ078_ko_amgS_frag1_F | gctatgaccatgattacggaatcgaccatgaactcccggtg |
| oSKJ079_ko_amgS_frag1_R | cggtccgatcgatgaaaacggcggcctgatacccgacg |
| oSKJ080_ko_amgS_frag2_F | cccgtcgggtatcaggccgccgttttcatcgatcggaccgg |
| oSKJ081_ko_amgS_frag2_R | catgcctgcaggtcgactctacctggccaagccgttcaac |
| oSKJ084_ko_PA1056_frag1_F | gctatgaccatgattacgcgttcggccacttcgagttg |
| oSKJ085_ko_PA1056_frag1_R | cgaacggctgatcatgcggcgatcagccaatggctcatggg |
| oSKJ086_ko_PA1056_frag2_F | ccatgagccattggctgatcgccgcatgatcagccgtt |
| oSKJ087_ko_PA1056_frag2_R | catgcctgcaggtcgactatctccttcagcggcgcttc |
| oSKJ090_ko_phoQ_frag1_F | gctatgaccatgattacggaataccaccacgacctggc |
| oSKJ091_ko_phoQ_frag1_R | gccaagtctcagactgtagcgatgcgcagggaacggatcac |
| oSKJ092_ko_phoQ_frag2_F | tgatccgttccctgcgcatcgctacagtctgagacttggcgg |
| oSKJ093_ko_phoQ_frag2_R | catgcctgcaggtcgactgcccacatttccgggaaacg |

**Table 4 Minimum inhibitory concentration (MIC) measurements of antibiotics**

| Clinically relevant antibiotics | MIC (μg/mL) |
|---|---|
| Tobramycin | 1 |
| Ciprofloxacin | 0.125 |
| Meropenem | 6 |

**Table 5 Supplementary data on transposon sequencing datasets**

Extended information on Tn-seq experimental data showing growth comparisons: liquid vs. liquid (UV light) and gel vs. liquid (UV liquid). The filename with growth can be found on Github.

i. Supplementary Tn-seq dataset showing growth under confinement (gel vs liquid (UV-treated))

ii. Supplementary Tn-seq dataset capturing mutant responses to tobramycin, comparing liquid versus liquid (UV light) and gel versus liquid (UV liquid); corresponding filename available on GitHub

iii. Supplementary Tn-seq dataset detailing antibiotic response comparisons, liquid versus liquid (UV light) and gel versus liquid (UV liquid) under colistin treatment, with the associated filename provided on GitHub.

iv. Gene ontology tables corresponding to the graphs are provided, covering multiple comparative conditions analyzed in the study.

All the above-mentioned files, including gene ontology tables and comparison datasets, are available on GitHub at the following link: https://github.com/monnappasourabh/antibiotic-tolerance.git
